# Muscle March5 restrains an ATF4–GDF15 endocrine stress axis that modulates feeding and body composition

**DOI:** 10.64898/2026.07.07.737120

**Authors:** Guixing Ma, Yong Chen, Siyuan Cheng, Yangshan Chen, Wei Pang, Litong Chen, Huiling Cao

**Author notes:** Address correspondence to: Dr. Huiling Cao. These authors contributed equally to this study (Guixing Ma, Yong Chen, Siyuan Cheng, Yangshan Chen).

## Abstract

Skeletal muscle can release endocrine stress signals during aging and wasting, but the upstream mechanisms that restrain this response remain incompletely defined. Here we identify March5 as a muscle proteostatic checkpoint that limits ATF4-dependent GDF15 production. March5 expression declined in aged and atrophic muscle, whereas muscle-specific March5 deletion induced ATF4 accumulation, marked GDF15 elevation, reduced food intake and progressive loss of body, muscle and bone mass. Restoration of feeding, GDF15 neutralization or muscle Atf4 deletion substantially attenuated the wasting phenotype. Mechanistically, March5 interacted with ATF4 and promoted its ubiquitination at K92, thereby limiting ATF4 stability and Gdf15 expression. Conversely, muscle March5 gain-of-function or pharmacological attenuation of ATF4 signaling improved feeding, body composition and physical performance in aged mice. These findings define a March5-ATF4-GDF15 endocrine stress axis linking muscle proteostatic control to feeding suppression and systemic body-composition remodeling.

## Introduction

Food intake is a fundamental behavior required for the maintenance of energy balance, body composition and organismal homeostasis^1^. It is coordinated by central neural circuits and peripheral endocrine signals that integrate information about nutrient availability, tissue stress, inflammation and energy demand^2–4^. Disruption of feeding behavior contributes to a broad spectrum of metabolic and degenerative conditions, including obesity, cachexia, anorexia of aging and sarcopenia^1,5,6^. In older individuals, reduced appetite and inadequate nutrient intake are common and are associated with frailty, loss of skeletal muscle mass, impaired mobility and increased morbidity^5,7–11^. Although many circulating factors, including leptin^12^, ghrelin^13^, GLP-1^14^ and growth differentiation factor 15 (GDF15) ^15–23^, have been implicated in the regulation of food intake, the tissue-derived mechanisms that link peripheral organ stress to altered feeding behavior remain incompletely understood^1^.

Skeletal muscle is increasingly recognized as an endocrine and metabolic organ that communicates with distant tissues through myokines, metabolites and stress-induced secreted factors^24^. During aging, disuse, inflammation and mitochondrial dysfunction, skeletal muscle undergoes profound remodeling that includes loss of contractile mass, altered substrate metabolism, impaired proteostasis and activation of cellular stress pathways^25–28^. These changes are not only cell-autonomous events within muscle but may also influence whole-body metabolism and feeding behavior. However, whether and how muscle-intrinsic proteostatic mechanisms restrain such endocrine stress outputs during aging or muscle wasting remains poorly defined.

GDF15 has emerged as a major stress-responsive hormone that suppresses food intake and regulates body weight through its hindbrain receptor GFRAL^17,18,22,29,30^. Elevated GDF15 is observed in diverse stress and disease contexts, including mitochondrial dysfunction, inflammation, cancer cachexia, tissue injury and aging^31^. In skeletal muscle, previous studies have shown that mitochondrial stress and integrated stress response activation can induce GDF15 expression, and muscle-derived GDF15 can contribute to anorexia and systemic metabolic remodeling under pathological stress conditions^32,33^. These observations establish GDF15 as an important endocrine output of stressed muscle. However, GDF15 biology is highly context dependent. GDF15 can participate in maladaptive anorexia and wasting, but it may also exert adaptive or protective effects in metabolic stress and aging-associated inflammation^16,31^. Therefore, defining the upstream mechanisms that control GDF15 production in specific tissues and disease contexts is essential for understanding when this pathway is beneficial or detrimental.

The integrated stress response transcription factor ATF4 is a key regulator of Gdf15 expression in response to mitochondrial stress, endoplasmic reticulum stress, nutrient deprivation and inflammatory stimuli^15,31,34^. In many settings, ATF4 activation is controlled through phosphorylation of eIF2α and preferential translation of ATF4 mRNA^35^. However, ATF4 abundance and activity can also be regulated at the level of protein stability, including through ubiquitin-dependent proteasomal degradation^27,36,37^. The molecular machinery that controls ATF4 protein stability in skeletal muscle, particularly under aging- or wasting-associated stress, remains incompletely understood. Identifying such regulatory mechanisms may reveal how muscle cells tune endocrine stress signaling without constitutively activating systemic appetite-suppressive pathways.

March5, also known as MARCHF5 or MITOL, is a membrane-associated RING-type E3 ubiquitin ligase localized primarily to mitochondria and other intracellular membranes^38^. March5 has been implicated in mitochondrial dynamics, organelle quality control, protein ubiquitination and cellular stress responses^39–43^. Because mitochondrial and proteostatic stress pathways are closely linked to ATF4 activation and GDF15 production, March5 represents a plausible upstream regulator of muscle endocrine stress signaling. However, whether March5 controls ATF4 stability in skeletal muscle and whether this regulation affects GDF15 production, feeding behavior and systemic body composition have not been established.

Here, we identify March5 as a muscle proteostatic checkpoint that restrains an ATF4-GDF15 endocrine stress program. We show that March5 expression declines in aged and atrophic skeletal muscle, whereas muscle-specific March5 deletion induces ATF4 accumulation, robust GDF15 elevation, reduced food intake and progressive loss of body, muscle and bone mass. Restoration of feeding, GDF15 neutralization and muscle Atf4 deletion substantially attenuate major components of the wasting phenotype. Mechanistically, March5 interacts with ATF4 and promotes ATF4 ubiquitination at lysine 92, thereby limiting ATF4 stability and Gdf15 expression. Conversely, March5 gain-of-function or pharmacological attenuation of ATF4-associated signaling improves feeding, body composition and physical performance in aged mice. These findings reposition March5 as a muscle proteostatic brake on ATF4-GDF15 endocrine stress signaling, providing a mechanism by which aged or wasting muscle can influence feeding and systemic body composition.

## Results

### March5 is reduced in aged and atrophic skeletal muscle

To identify muscle-intrinsic regulators associated with aging- and wasting-related muscle remodeling, we first examined March5, a membrane-associated E3 ubiquitin ligase implicated in organelle proteostasis and cellular stress responses. Exploratory analyses of public genetic and protein-interaction datasets suggested that March5 and March5-associated proteins were linked to muscle and musculoskeletal phenotypes (Extended Data Fig. 1a, b and Supplementary Table 1). We next examined March5 expression in mouse and human skeletal muscle.

Immunohistochemical analysis showed that March5 protein abundance was reduced in skeletal muscle from aged mice compared with young controls (Fig. 1a, b). A similar reduction in March5 staining was observed in skeletal muscle samples from older human individuals compared with younger individuals (Fig. 1c, d and Supplementary Table 2). In cultured C2C12 myogenic cells, oxidative stress induced by hydrogen peroxide decreased March5 protein levels (Fig. 1e, f). Inflammatory stimulation with IL-1β also reduced March5 protein abundance (Fig. 1g, h). Consistently, March5 expression was decreased in immobilized skeletal muscle, a model of disuse-associated muscle atrophy (Fig. 1i–l).

**Figure 1.**
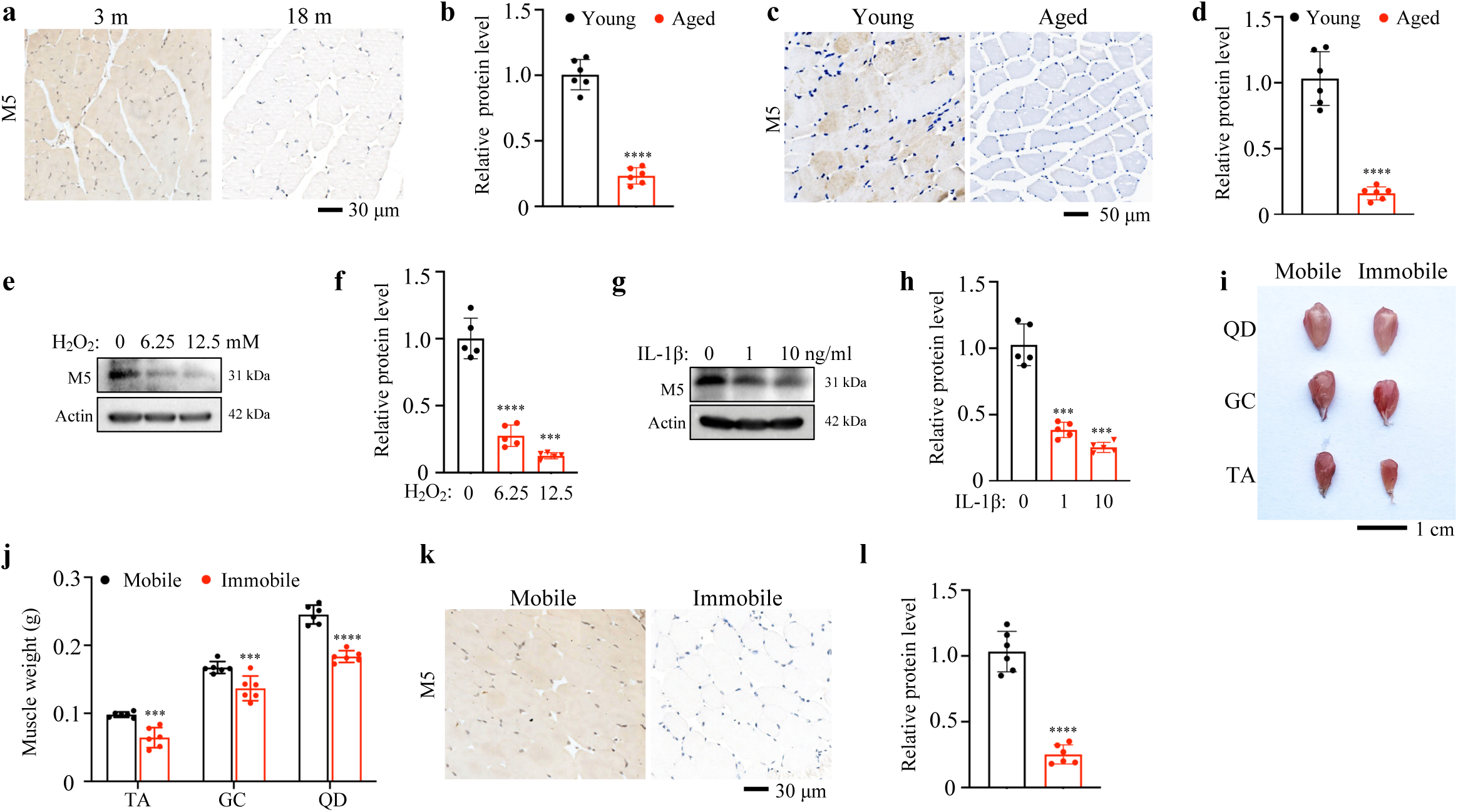
March5 expression level is associated with muscle function. (**a-d**) IHC staining. Quadriceps (QD) sections from 3- and 18-month-old male C57/BL6 mice (**a**) and skeletal muscle sections from young and aged male human individuals (**c**) were subjected to IHC staining using an anti-March5 antibody. Quantification (**b and d**) of IHC staining results shown in (**a**) and (**c**), respectively. N = 6 mice per group in (**a**). N = 6 individuals per group in (**c**). Scale bars, 30 μm in (**a**) and 50 μm in (**c**). (**e-h**) WB analyses. Protein extracts from C2C12 cells after H_2_O_2_ (**e**) or IL-1β treatment **(g**) for 72 hours were analyzed via WB. (**f**) Quantification of (**e**). (**h**) Quantification of (**g**). (**i and j**) Representative images (**i)** and statistical analysis (**j)** of quadriceps (QD), gastrocnemius (GC), and tibialis anterior (TA) muscles following 14 days of hindlimb immobilization. Male mice, N = 6 per group. Scale bar, 1 cm. (**k**) IHC staining. Skeletal muscle sections from immobilized and control mice were subjected to IHC staining using an anti-March5 antibody. Male mice, N = 6 per group. Scale bar, 30 μm. (**l**) Quantification of (**k**). *P < 0.05, **P < 0.01, ***P < 0.001, ****P < 0.0001, versus controls. Student’s *t* test. Results are expressed as the mean ± standard deviation (sd).

These data show that March5 is reduced in aged, oxidative, inflammatory and atrophic muscle contexts. We therefore generated skeletal muscle-specific March5 knockout mice to determine the in vivo consequences of March5 loss in muscle.

### Muscle-specific March5 deletion causes progressive post-weaning wasting

To determine the *in vivo* consequence of March5 loss in muscle, we crossed March5-floxed mice with CKMM-Cre mice to generate conditional March5 knockout mice (hereafter referred to as cKO mice). cKO mice were born at the expected Mendelian frequency and showed no gross abnormalities at birth. Alizarin red and Alcian blue staining at postnatal day 0 revealed no detectable differences in skeletal patterning, mineralization or bone length between cKO mice and littermate controls (Extended Data Fig. 2a–c).

At one month of age, cKO mice showed comparable body size, hindlimb muscle mass, forelimb grip strength, trabecular bone parameters and adipose tissue mass relative to controls (Extended Data Fig. 2d–l). March5 transcript analysis across the tissues examined confirmed efficient reduction in skeletal muscle (Extended Data Fig. 2m).

After weaning, body weight progressively diverged between cKO and control mice in both sexes (Fig. 2a,b). Male cKO mice developed premature mortality beginning at approximately 40 weeks of age, whereas female cKO mice did not show a comparable survival phenotype during the period examined (Fig. 2c).

**Figure 2.**
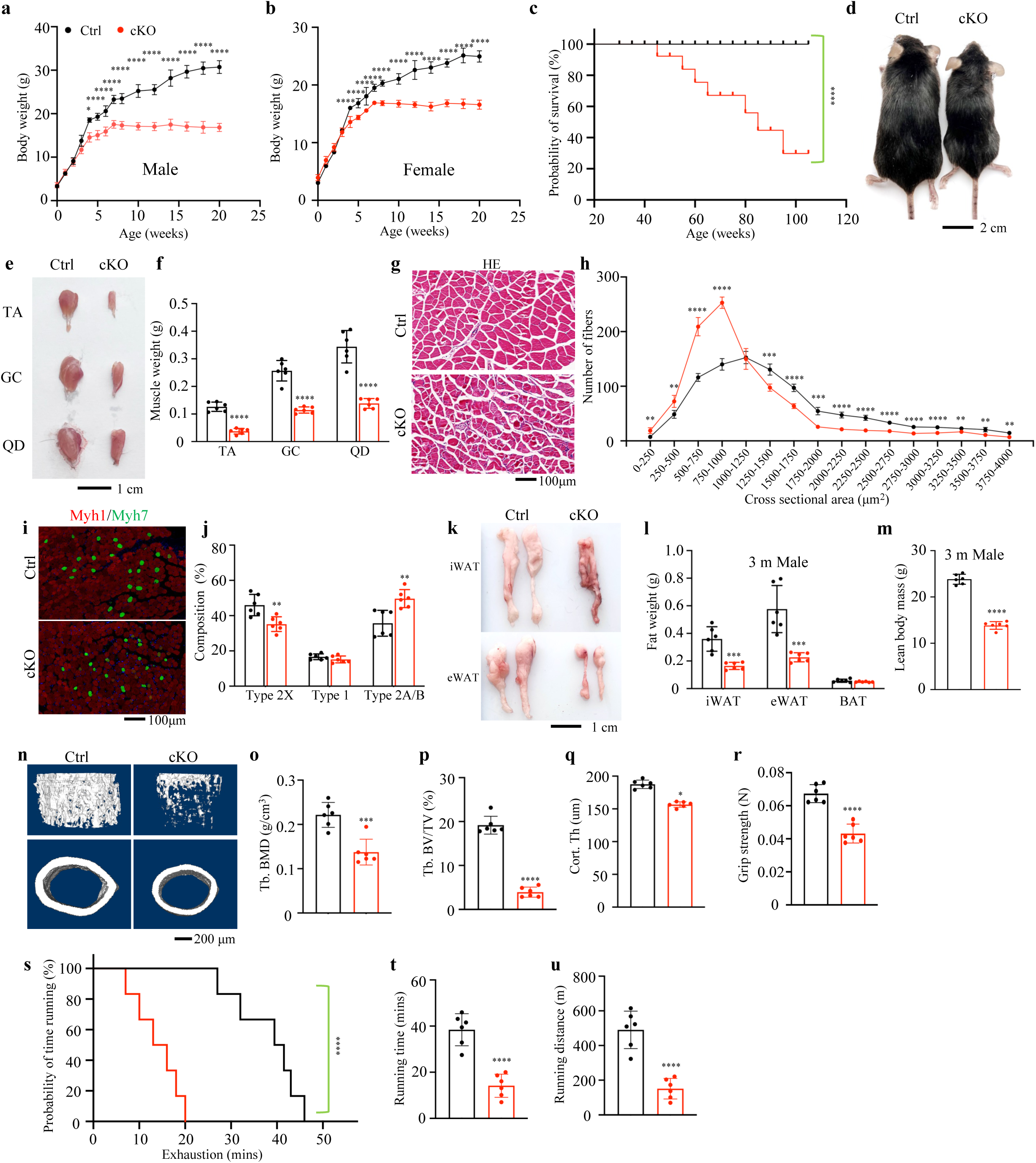
Muscle-specific March5 deletion causes progressive post-weaning wasting. (**a-c**) Body weight measurements of male (**a)** and female **(b**) mice, and survival curve (**c**) of male mice with skeletal muscle-specific March5 deficiency (cKO) compared to littermate controls. (**d**) Representative images of cKO mice and littermate controls. Scale bar, 2 cm. (**e and f**) Representative images (**e**) and statistical analysis (**f**) of TA, GC, and QD muscles from 3-month-old cKO mice and littermate controls. Male mice, N = 6 for per group. Scale bar, 1 cm. (**g**) HE staining of skeletal muscle sections from cKO and control mice. Male mice, N = 6 per group. Scale bar, 100 μm. (**h**) Quantification of muscle fiber diameter based on (**g**). (**i**) IF staining of extensor digitorum longus (EDL) muscle sections from cKO and littermate control mice using antibodies against Myh1 (marker of fast type 2X fibers) and Myh7 (marker for slow type 1 fibers). Male mice, N = 6 per group. Scale bar, 100 μm. (**j**) Quantification of (**i**). (**k and l**) Representative images (k) and quantification (l) of BAT, iWAT, and eWAT in cKO and control mice at 3 months of age. Scale bar, 1 cm. (**m**) Lean body mass quantification in cKO and control mice at 3 months of age. N = 6 mice per group. (**n-q**) 3D reconstruction (**n**) and quantitative analyses of BMD (**o**), BV/TV (**p**), and Cort.Th (**q**) from μCT scans of distal femurs in 3-month-old cKO and control mice. Scale bar, 200 μm. N = 6 mice per group. (**r-u**) Evaluation of exercise capacity in mice. Quantitative analyses of forelimb grip strength (**r**), treadmill running time (**s and t**), and running distance (**u**). *P < 0.05, **P < 0.01, ***P < 0.001, ****P < 0.0001, versus controls. Student’s *t* test. Results are expressed as the mean ± standard deviation (sd).

At three months of age, cKO mice displayed a visibly smaller body size and reduced hindlimb muscle mass (Fig. 2d–f). The weights of tibialis anterior, gastrocnemius and quadriceps muscles were lower in cKO mice than in controls (Fig. 2e,f). Haematoxylin and eosin staining showed a shift toward smaller myofibre cross-sectional areas in cKO muscle (Fig. 2g,h). Immunofluorescence staining of extensor digitorum longus sections with Myh1 and Myh7 antibodies revealed a reduction in Myh1-positive fibres and changes in the remaining fibre population (Fig. 2i,j). Because Myh2 and Myh4 were not directly assessed in this experiment, Myh1-negative/Myh7-negative fibres were not assigned to specific type 2A or type 2B subtypes.

The cKO phenotype extended to other components of body composition. Lean body mass and white adipose tissue mass were reduced in cKO mice (Fig. 2k–m). Micro-computed tomography of distal femurs showed lower trabecular bone mineral density, trabecular bone volume fraction and cortical thickness in cKO mice (Fig. 2n–q). Functional assays showed lower forelimb grip strength, shorter treadmill running time and reduced running distance (Fig. 2r–u). Echocardiographic parameters and peripheral oxygen saturation did not differ significantly between genotypes at two months of age (Extended Data Fig. 3a–i).

Together, these data show that CKMM-Cre-driven March5 deletion causes progressive post-weaning loss of body mass, muscle mass, adipose tissue mass and bone mass, accompanied by impaired physical performance.

### Muscle March5 gain-of-function produces reciprocal musculoskeletal phenotypes

To complement the loss-of-function analysis, we generated transgenic mice overexpressing March5 in skeletal muscle (hereafter referred to as TG mice). At three months of age, TG mice showed a modest but significant increase in body weight compared with littermate controls (Fig. 3a,b). Gross dissection showed increased mass of gastrocnemius and tibialis anterior muscles in TG mice, whereas quadriceps mass showed no significant increase under the conditions examined (Fig. 3c,d). Histological analysis revealed larger myofibre cross-sectional areas in TG muscle (Fig. 3e,f). Myh1 and Myh7 staining further showed altered fibre composition in TG mice compared with controls (Fig. 3g,h). Micro-computed tomography showed higher trabecular bone mineral density and trabecular bone volume fraction in TG mice than in controls (Fig. 3i–k). TG mice also exhibited higher forelimb grip strength, longer treadmill running time and increased running distance (Fig. 3l–o).

**Figure 3.**
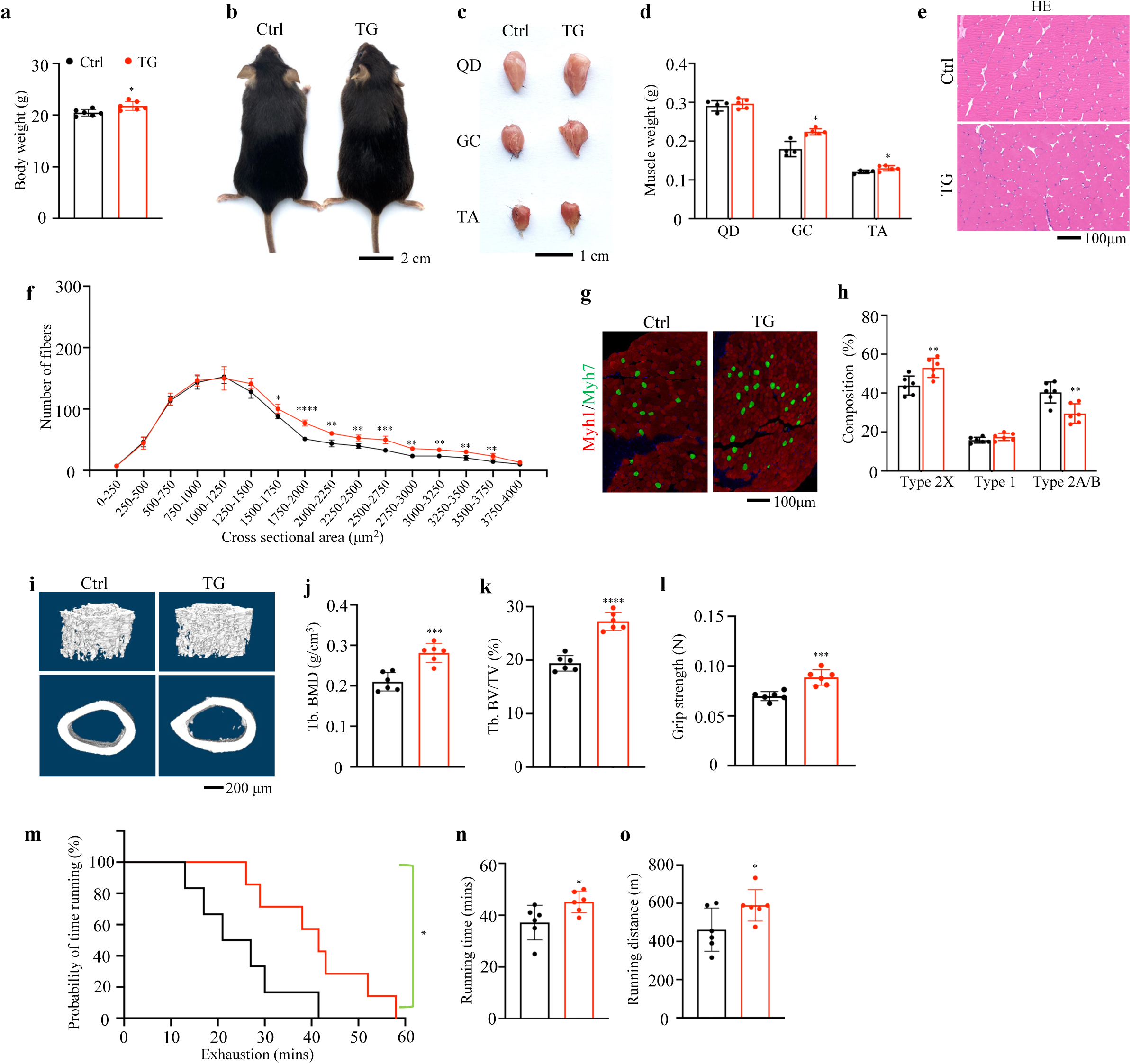
March5 overexpression in skeletal muscle enhances musculoskeletal function in mice. (**a**) Body weight measurements of mice with skeletal muscle specific March5 overexpression (TG) and littermate controls. (**b**) Representative images of TG mice and littermate controls. Scale bar, 2 cm. (**c and d**) Representative images and statistical analysis of QD, GC, TA from 3-month-old TG mice and littermate controls. Male mice, N = 6 mice per group. Scale bar, 1 cm. (**e**) HE staining of skeletal muscle sections from TG and control mice. Male mice, N = 6 per group. Scale bar, 100 μm. (**f**) Quantification of muscle fiber diameter based on (**e**). (**g**) IF staining of extensor digitorum longus (EDL) muscle sections from TG and littermate control mice using antibodies against Myh1 (marker of fast type 2X fibers) and Myh7 (marker for slow type 1 fibers). Male mice, N = 6 per group. Scale bar, 100 μm. (**h**) Quantification of (**g**). (**i-k**) 3D reconstruction (**i**) and quantitative analyses of BMD (**j**) and BV/TV (**k**) from μCT scans of distal femurs in 3-month-old TG and control mice. Scale bar, 200 μm. N = 6 mice per group. (**l-o**) Evaluation of exercise capacity in mice. Quantitative analyses of forelimb grip strength (**l**), treadmill running time (**m-n**), and running distance (**o**). *P < 0.05, **P < 0.01, ***P < 0.001, ****P < 0.0001, versus controls. Student’s *t* test. Results are expressed as the mean ± standard deviation (sd).

Thus, March5 gain-of-function in muscle produced reciprocal changes in muscle mass, bone parameters and physical performance relative to March5 loss.

### March5 deficiency induces a GDF15-associated endocrine stress signature

To define molecular changes associated with March5 deficiency, we performed RNA sequencing of skeletal muscle from three-month-old cKO and control mice. Differential-expression analysis revealed extensive transcriptional remodelling in cKO muscle (Fig. 4a,b). Pathway enrichment analysis identified gene sets related to muscle contraction, glycogen metabolism, mitochondrial organization, ion homeostasis, hypoxia responses, lipid catabolism and hormone responses (Fig. 4a).

**Figure 4.**
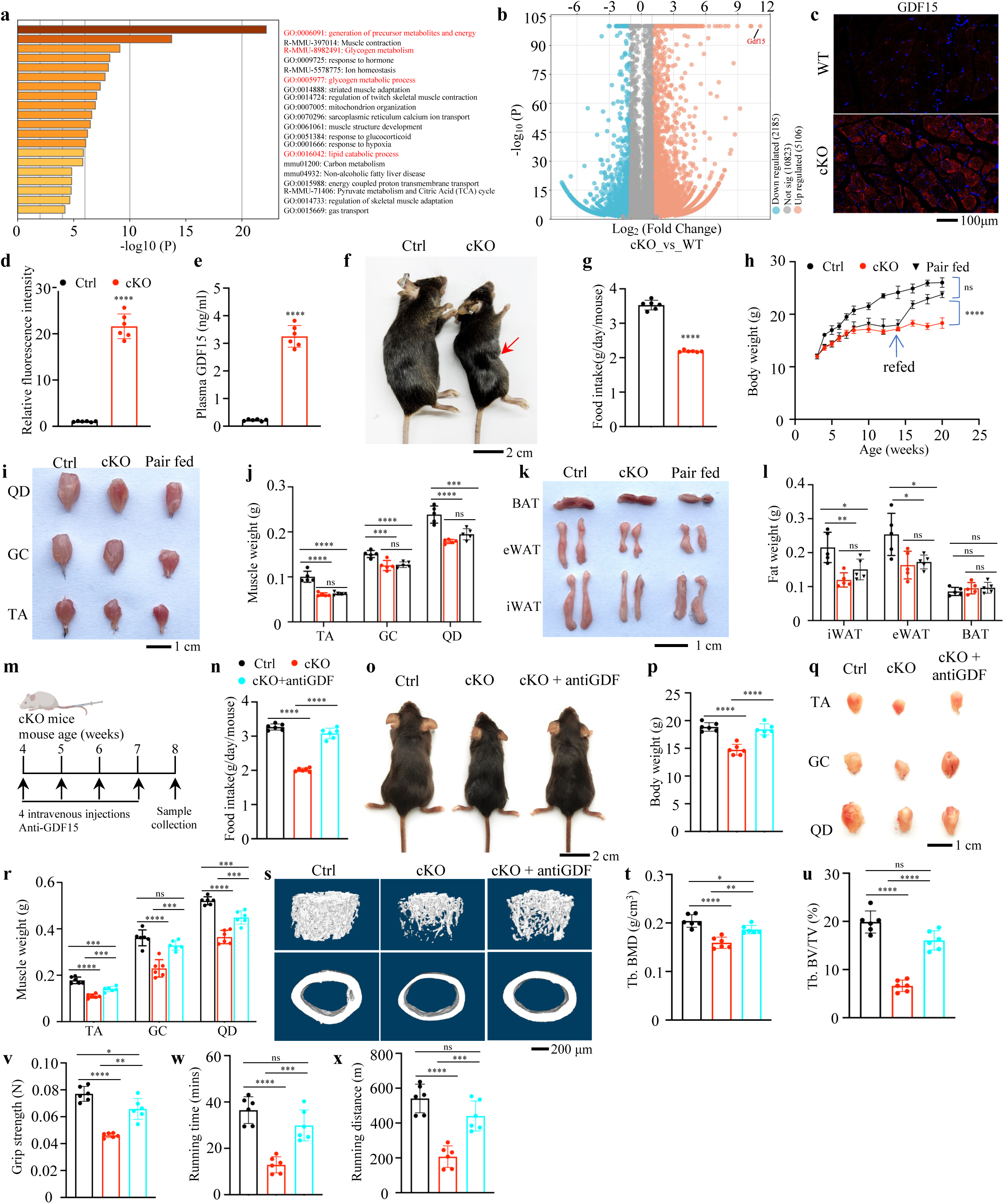
March5 knockout in skeletal muscle induces post-weaning wasting through GDF15-mediated food intake. (**a-b**) Heat map (**a**) and volcano plot (**b**) depicting differentially expressed genes in skeletal muscles of cKO mice compared to control mice at 3 months of age. (**c**) IF staining of QD sections from cKO and control mice using an anti-GDF15 antibody. Male mice, N = 6 per group. Scale bar, 100 μm. (**d**) Quantification of (**c**). (**e**) ELISA quantification of serum GDF15 levels in cKO and control mice. Male mice, N = 6 per group. (**f**) Representative images of cKO and control mice. The red arrow indicates the the abdominal area. Scale bar, 2 cm. (**g**) Food intake statistics of cKO and control mice at 3 months of age. (**h**) Body weight of control (Ctrl) and muscle-specific March5 knockout (cKO) mice under pair-feeding and refeeding conditions. (**i-j**) Representative images and quantification of skeletal muscle from Ctrl and cKO mice under pair-feeding. Scale bar, 1 cm. (**k-l**) Representative images and quantification of adipose tissue from Ctrl and cKO mice under pair-feeding. Scale bar, 1 cm. Male mice, N = 5 per group. (**m**) Schematic diagram of experimental design for GDF15 neutralization using an anti-GDF15 antibody. (**n**) Food intake statistics of cKO mice, cKO + ant-GDF antibody treatment group, and control mice. (**o and p**) Representative images (o) and body weight (p) of cKO mice, cKO + antiGDF mice, and control mice. Scale bar, 2 cm. (**q** and **r**) Representative images (q) and statistical analysis (r) of QD, GC, and TA from cKO, cKO + antiGDF, and control mice. Male mice, N = 6 per group. Scale bar, 1 cm. (**s-u**) 3D reconstruction (**s**) and quantitative analyses of BMD (**t**) and BV/TV (**u**) from μCT scans of distal femurs in cKO, cKO+antiGDF, and control mice. Scale bar, 200 μm. N = 6 mice per group. (**v-x**) Evaluation of exercise capacity in mice. Quantitative analyses of forelimb grip strength (**v**), treadmill running time (**w**), and running distance (**x**). *P < 0.05, **P < 0.01, ***P < 0.001, ****P < 0.0001, versus controls. Student’s *t* test. Results are expressed as the mean ± standard deviation (sd).

Among the most strongly induced transcripts, Gdf15, a key molecule involved in dietary regulation, stress response, inflammation, and systemic metabolism ^15,17,21,22,30,44–46^, showed an increase of more than 2,000-fold in cKO muscle (Fig. 4b). Immunofluorescence staining confirmed increased GDF15 protein abundance in quadriceps sections from cKO mice (Fig. 4c,d). ELISA analysis showed that circulating GDF15 concentrations were markedly higher in cKO mice than in controls (Fig. 4e).

Food intake was lower in cKO mice than in control mice at three months of age (Fig. 4f,g). Reduced food intake was accompanied by lower lean body mass and reduced inguinal and epididymal white adipose tissue weights (Fig. 2kl). Respiratory exchange ratio and energy expenditure did not differ significantly between genotypes in the metabolic-cage assays performed, and circulating thyroxine, corticosterone and leptin concentrations were comparable (Extended Data Fig. 4a–h).

To assess the contribution of reduced food intake to the body-composition phenotype, control mice were pair-fed to match the food intake of cKO mice. During the restriction phase, pair-fed control mice lost body weight to an extent similar to cKO mice (Fig. 4h). After ad libitum feeding was restored, pair-fed control mice regained body weight, whereas cKO mice remained lighter than controls (Fig. 4h). Skeletal muscle and white adipose tissue weights showed corresponding changes across the feeding and refeeding phases (Fig. 4i–l).

We next examined the functional contribution of GDF15 by administering a GDF15-neutralizing antibody^30,46^ to cKO mice once weekly for four weeks (Fig. 4m). Antibody-treated cKO mice showed increased food intake and body weight compared with IgG-treated cKO mice (Fig. 4n–p). GDF15 neutralization increased tibialis anterior, gastrocnemius and quadriceps muscle weights (Fig. 4q,r), improved trabecular bone mineral density and bone volume fraction (Fig. 4s–u), and increased grip strength, treadmill running time and running distance (Fig. 4v–x).

These data identify GDF15 as a prominent endocrine factor induced by March5 deficiency and show that reduced food intake and GDF15 signalling contribute to the systemic phenotype of cKO mice.

### ATF4 is required for GDF15 induction and systemic phenotypes following March5 loss

Because Gdf15 is transcriptionally responsive to ATF4^1,15,295,37,49^, we examined ATF4 abundance in March5-deficient muscle. Immunohistochemistry and immunoblotting showed higher ATF4 protein levels in cKO skeletal muscle than in control muscle (Fig. 5a,b and Extended Data Fig. 5a,b). In C2C12 cells, March5 knockdown did not significantly increase Atf4 or Eif2a mRNA abundance, but it markedly increased Gdf15 mRNA abundance (Fig. 5c). At the protein level, March5 knockdown increased ATF4 and GDF15 without increasing total eIF2α^47^ (Fig. 5d,e). Conversely, expression of wild-type March5 reduced ATF4 and GDF15 protein abundance, whereas the ligase-impaired mutant M5 (M5^H43W^)^48^ did not reproduce this effect (Fig. 5f,g). March5 manipulation did not appreciably alter total eIF2αabundance under these conditions.

**Figure 5.**
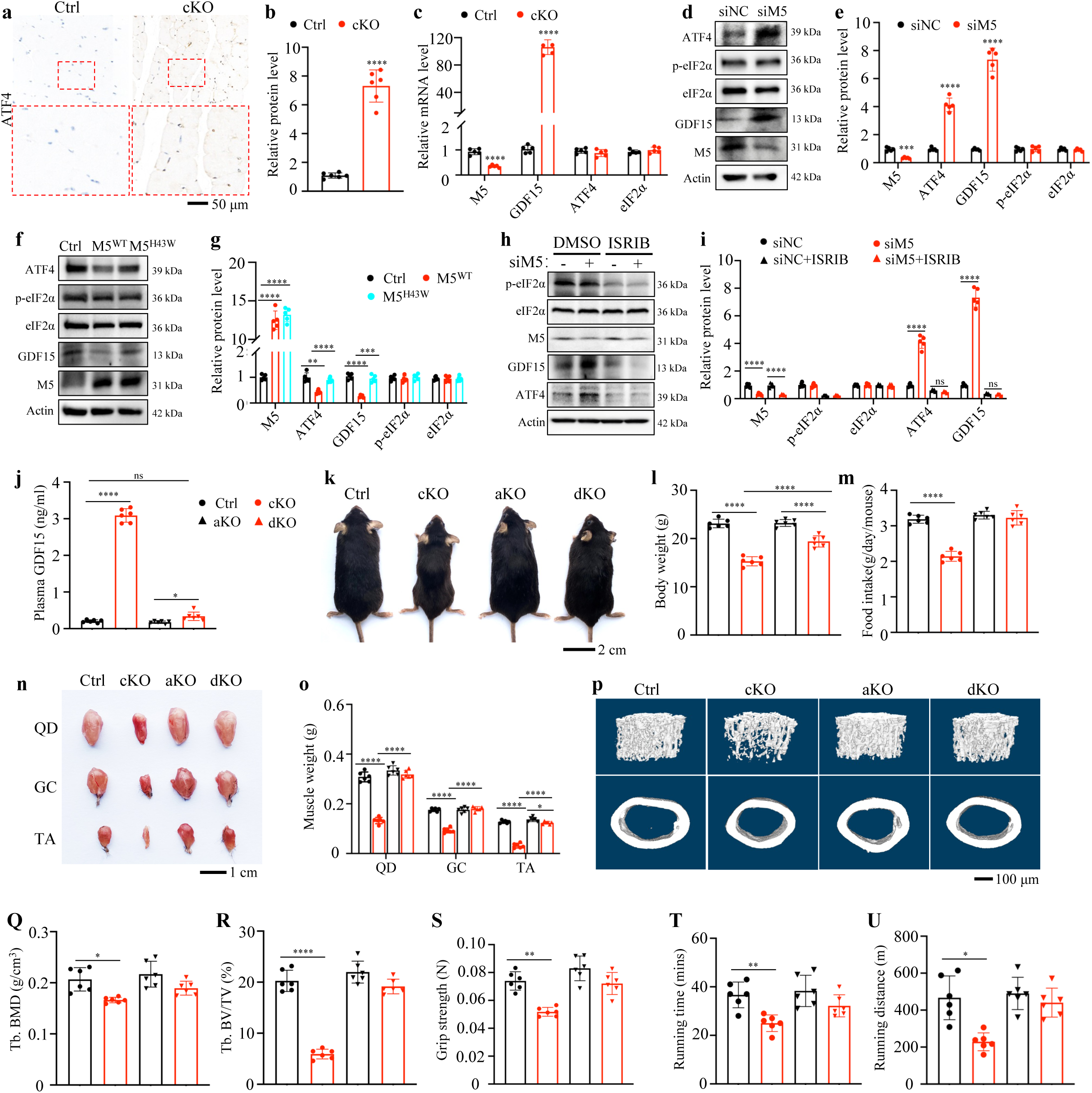
ATF4 Upregulation Mediates GDF15 Elevation in March5 Knockout Mice. (**a**) IHC staining of skeletal muscle sections from cKO and control mice using an anti-ATF4 antibody. Male mice, N = 6 per group. Scale bar, 50 μm. (b) Quantification of (**a**). (**c**) Quantitative real-time PCR (qRT-PCR) analysis of RNA isolated from QD muscles from cKO and control mice. (**d**) WB analyses of protein extracts from C2C12 cells following March5 knockdown (siM5). (**e**) Quantification of (**d**). (**f**) WB analyses of protein extracts from C2C12 cells overexpressing WT (M5^WT^) or the H43W mutant (M5^H43W^). (**g**) Quantification of (**f**). (**h**) WB analyses of protein extracts from C2C12 cells after March5 knockdown (siM5) or ISRIB treatment. (**i**) Quantification of (**h**). (**j**) ELISA quantification of serum GDF15 levels in cKO, aKO, dKO and control mice. Male mice, N = 6 per group. (**k and l**) Representative images (**k**) and Body weight (**l**) of cKO, ATF knockout (aKO), March5 and ATF4 double knockout (dKO) mice, and control mice. Scale bar, 2 cm. (**m**) Food intake statistics for cKO, aKO, dKO, and control mice. (**n and o**) Representative images (**n)** and statistics analysis (**o)** of QD, GC, and TA muscles in the 4 groups. Male mice, N = 6 per group. Scale bar, 1 cm. (**p-r**) 3D reconstruction (**p**) and quantitative analyses of BMD (**q**) and BV/TV (**r**) from μCT scans of distal femurs of mice in the 4 groups. Scale bar, 200 μm. N = 6 mice per group. (**s-u**) Evaluation of exercise capacity in mice, including forelimb grip strength (**s**), treadmill running time (**t**), and running distance (**u**). *P < 0.05, **P < 0.01, ***P < 0.001, ****P < 0.0001, versus controls. Student’s *t* test. Results are expressed as the mean ± standard deviation (sd).

ISRIB treatment reduced ATF4 and GDF15 accumulation after March5 knockdown in C2C12 cells (Fig. 5h,i). To test the requirement for ATF4 in *vivo*, we generated mice with muscle-targeted deletion of Atf4 alone or combined deletion of March5 and Atf4. Deletion of Atf4 reduced circulating GDF15 concentrations in March5-deficient mice (Fig. 5j). Compared with cKO mice, double-knockout mice consumed more food and had higher body weight, although body weight remained below that of control mice (Fig. 5k–m).

Double-knockout mice also had higher tibialis anterior, gastrocnemius and quadriceps muscle weights than cKO mice (Fig. 5n,o). Trabecular bone mineral density and bone volume fraction were increased relative to cKO mice (Fig. 5p–r). Grip strength, treadmill running time and running distance were also improved in double-knockout mice compared with cKO mice (Fig. 5s–u). These data show that ATF4 is required for the GDF15 increase and for multiple systemic phenotypes caused by March5 loss.

### March5 promotes ATF4 ubiquitination through a ATF4^K92^-dependent mechanism

We next explored the molecular mechanism by which March5 regulates ATF4. The increase in ATF4 protein without a corresponding increase in Atf4 mRNA suggested post-transcriptional regulation. Co-immunoprecipitation assays showed an association between March5 and ATF4 in C2C12 cells (Fig. 6a). Immunofluorescence analysis showed partial co-localization of the two proteins (Fig. 6b).

**Figure 6.**
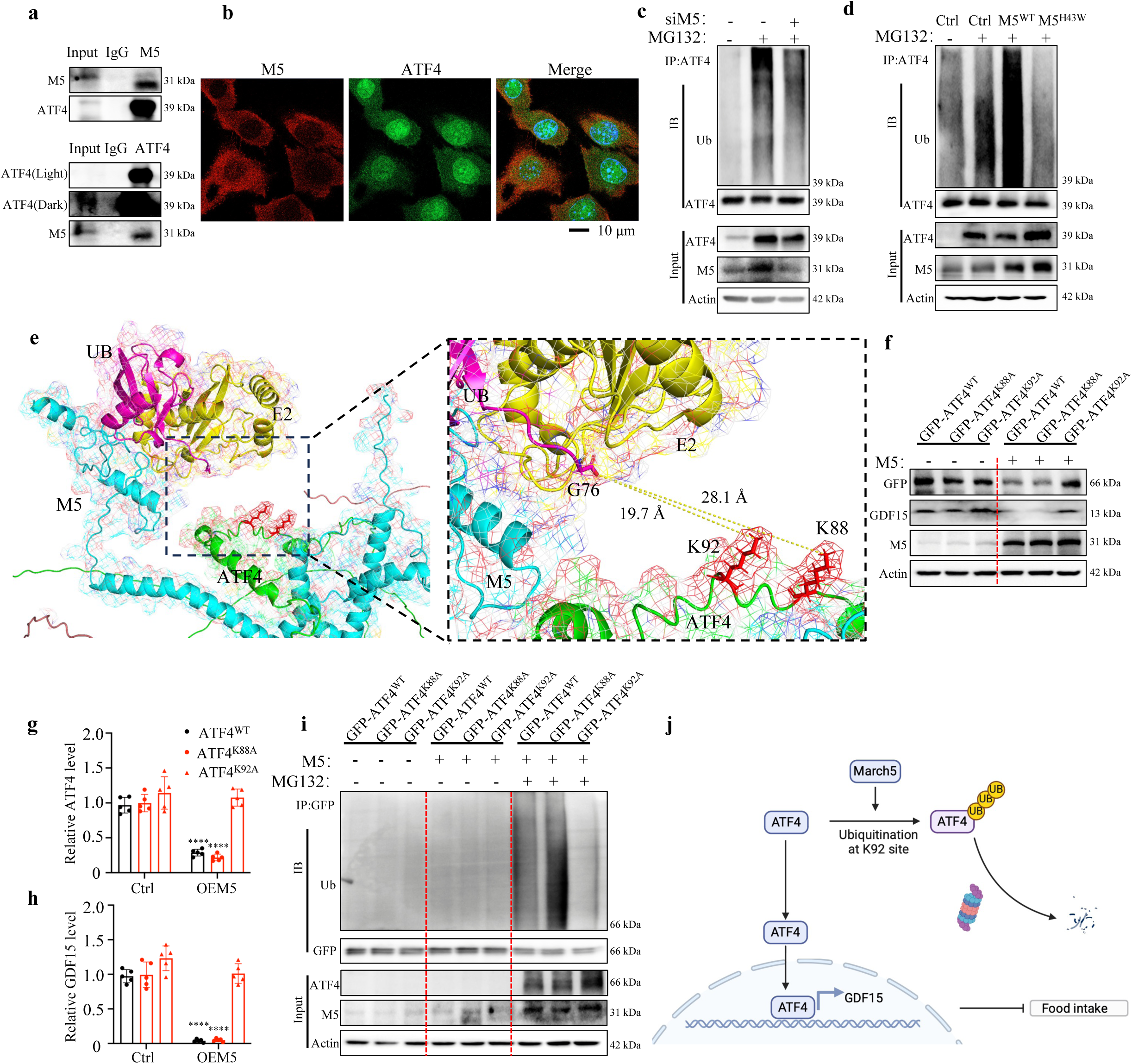
March5 modulates GDF15 expression through direct interaction and ubiquitination of ATF4. (**a**) Co-immunoprecipitation (Co-IP) assays. Cell lysates of C2C12 cells were subjected to immunoprecipitation (IP) and immunoblotting (IB) to detect the endogenous interaction between ATF4 and March5. (**b**) IF staining of C2C12 cells using antibodies against March5 and ATF4, demonstrating their colocalization. Scale bar, 10 μm. (**c**) WB analyses of protein extracts from C2C12 cells following MG132 treatment and March5 knockdown, using the indicated antibodies. (**d**) WB analyses of protein extracts from C2C12 cells overexpressing M5^WT^ or M5^H43W^ after MG132 treatment, using the indicated antibodies. (**e**) Predicted protein structure of the March5-ATF4-E2-UB complex using AlphaFold-multimer, visualized with PyMOL. Ubiquitins (UBs) are shown in magenta, E2 in orange, March5 in cyans, and potential ubiquitination sites on ATF4 (K88 and K92) in red. (**f**) WB analyses of protein extracts from C2C12 cells overexpressing March5 (M5), ATF4 (GFP), or ATF4 mutants following MG132 treatment, using the indicated antibodies. (**g** and **h**) Quantification of ATF4 (g) and GDF15 (h) levels based on (F). (**i**) WB analyses of protein extracts from C2C12 cells overexpressing M5, ATF4, or ATF4 mutants after MG132 treatment, using the indicated antibodies. (**j**) Schematic diagram illustrating the mechanism by which March5 regulates food intake via modulation ATF4 protein stability. *P < 0.05, **P < 0.01, ***P < 0.001, ****P < 0.0001, versus controls. Student’s *t* test. Results are expressed as the mean ± standard deviation (sd).

We next examined ATF4 ubiquitination. March5 knockdown decreased ATF4 ubiquitination in MG132-treated cells (Fig. 6c). Conversely, expression of wild-type March5 increased ATF4 ubiquitination, whereas the ligase-impaired M5^H43W^ mutant did not (Fig. 6d). These data indicate that March5 ligase activity is required for efficient ATF4 ubiquitination.

To identify candidate March5-responsive lysines in ATF4, we used AlphaFold-Multimer^49,50^ to generate a structural model of a March5–E2–ubiquitin–ATF4 complex. In this model, the C-terminal G76 residue of ubiquitin was positioned near the N-terminal region of ATF4, with K92 and K88 located approximately 19.7 Å and 28.1 Å away, respectively (Fig. 6e). We therefore generated K88A and K92A ATF4 mutants.

Wild-type March5 reduced the abundance of wild-type ATF4 and the downstream product GDF15 (Fig. 6f–h). The K92A substitution markedly attenuated the effect of March5 on ATF4 and GDF15 abundance, whereas the K88A substitution had a weaker effect (Fig. 6f–h). In ubiquitination assays, March5-dependent ubiquitination was reduced for ATF4 K92A compared with wild-type ATF4 (Fig. 6i). These data identify K92 as a critical residue for March5-dependent regulation of ATF4 ubiquitination and turnover.

In summary, these findings indicate that March5 suppresses GDF15 expression by directly interacting with ATF4 and promoting its ubiquitination at the K92 site (Fig. 6j).

### Modulation of the March5–ATF4 axis improves aging-associated phenotypes

We next examined the March5–ATF4–GDF15 axis in aged muscle. ATF4 and GDF15 immunoreactivity was higher in skeletal muscle specimens from older human donors than in samples from younger donors (Fig. 7a–d).

**Figure 7.**
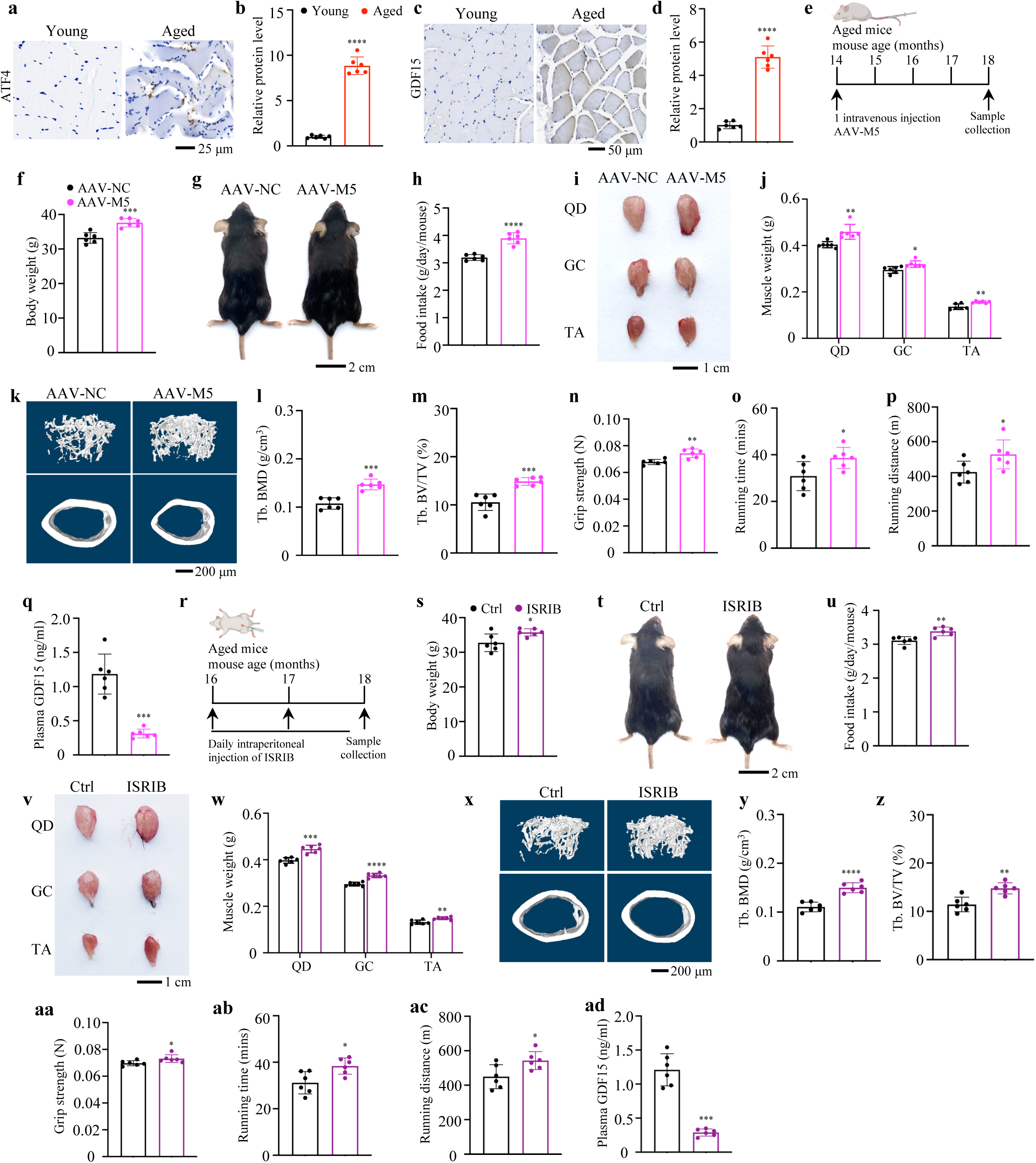
Modulation of the March5–ATF4 axis improves aging-associated phenotypes. (**a-d**) IHC staining of skeletal muscle sections from young and aged human individuals using antibodies against ATF4 (**a**) or GDF15 (**c**). (**b and d**) Quantifications based on (**a)** and **(c**). Male, N = 6 individuals per group. Scale bar, 25 and 50 μm. (**e**) Schematic diagram illustrating the experimental design for AAV-M5 or AAV-NC injections. (**f and g**) Representative images (**f**) and body weight (**g**) of mice treated with AAV-M5 or AAV-NC. Scale bar, 2 cm. (**h**) Food intake statistics for mice treated with AAV-M5 or AAV-NC. (**i and j**) Representative images (**i**) and statistics analysis (**j**) of QD, GC, and TA muscles in AAV-M5 or AAV-NC treated mice. Male mice, N = 6 per group. Scale bar, 1 cm. (**k-m**) 3D reconstruction (**k**) and quantitative analyses of BMD (**l**) and BV/TV (**m**) from μCT scans of distal femurs in AAV-M5 or AAV-NC treated mice. Scale bar, 200 μm. N = 6 mice per group. (**n-p**) Assessment of exercise capacity in mice, including forelimb grip strength (**n**), treadmill running time (**o**), and running distance (**p**). (**q**) ELISA quantifications of serum GDF15 levels in mice treated with AAV-M5 or AAV-NC. Male mice, N = 6 per group. (**r**) Schematic diagram describing the experimental design for ISRIB treatment. (**s and t**) Representative images (**s)** and body weight (**t**) of ISRIB treated or control mice. Scale bar, 2 cm. (**u**) Food intake statistics of ISRIB-treated or control mice. (**v and w**) Representative images (**v)** and statistics analysis (**w)** of QD, GC, and TA muscles in ISRIB treated or control mice. Male mice, N = 6 per group. Scale bar, 1 cm. (**x-z**) 3D reconstruction (**x**) and quantitative analyses of BMD (**y**) and BV/TV (**z**) from μCT scans of distal femurs of ISRIB treated or control mice. Scale bar, 200 μm. N = 6 mice per group. (**aa-ac**) Assessment of exercise capacity in mice, including forelimb grip strength (**aa**), treadmill running time (**ab**), and running distance (**ac**). (**ad**) ELISA assays measuring serum GDF15 in ISRIB-treated or control mice. N = 6 male mice per group. *P < 0.05, **P < 0.01, ***P < 0.001, ****P < 0.0001, versus controls. Student’s *t* test. Results are expressed as the mean ± standard deviation (sd).

To test whether increasing March5 abundance alters aging-associated phenotypes, 14-month-old male mice received an intravenous injection of AAV9 expressing March5 under the Acta1 promoter or a control vector and were analysed four months later (Fig. 7e, Extended Data Fig. 6). At approximately 18 months of age, AAV–March5-treated mice had higher body weight and food intake than control-vector-treated mice (Fig. 7f–h). Tibialis anterior, gastrocnemius and quadriceps muscle weights were also increased after AAV–March5 treatment (Fig. 7i,j). AAV–March5 treatment increased trabecular bone mineral density and bone volume fraction (Fig. 7k–m), and improved grip strength, treadmill running time and running distance (Fig. 7n–p). These changes were accompanied by lower circulating GDF15 concentrations (Fig. 7q).

As a complementary intervention, 16-month-old male mice were treated daily with ISRIB or vehicle for two months and analysed at approximately 18 months of age (Fig. 7r). ISRIB-treated mice had higher body weight and food intake than vehicle-treated mice (Fig. 7s–u). ISRIB treatment increased hindlimb muscle weights (Fig. 7v,w), trabecular bone mineral density and bone volume fraction (Fig. 7x–z), and improved grip strength and treadmill performance (Fig. 7aa–ac). Circulating GDF15 concentrations were lower in ISRIB-treated mice (Fig. 7ad). Histological examination of major organs did not reveal overt morphological abnormalities after ISRIB treatment (Extended Data Fig. 7).

These experiments show that increasing March5 abundance or attenuating ATF4-associated signalling modifies feeding, body composition and physical performance in aged mice.

### March5-dependent effects on feeding are retained in leptin-deficient mice

To determine whether the effects associated with March5 deficiency were retained in a hyperphagic model, we crossed cKO mice with leptin-deficient ob/ob mice. As expected, ob/ob mice showed higher food intake and body weight than control mice (Fig. 8a–c). Food intake was lower in cKO;ob/ob mice than in ob/ob mice, and body weight was correspondingly reduced (Fig. 8a–c).

**Figure 8.**
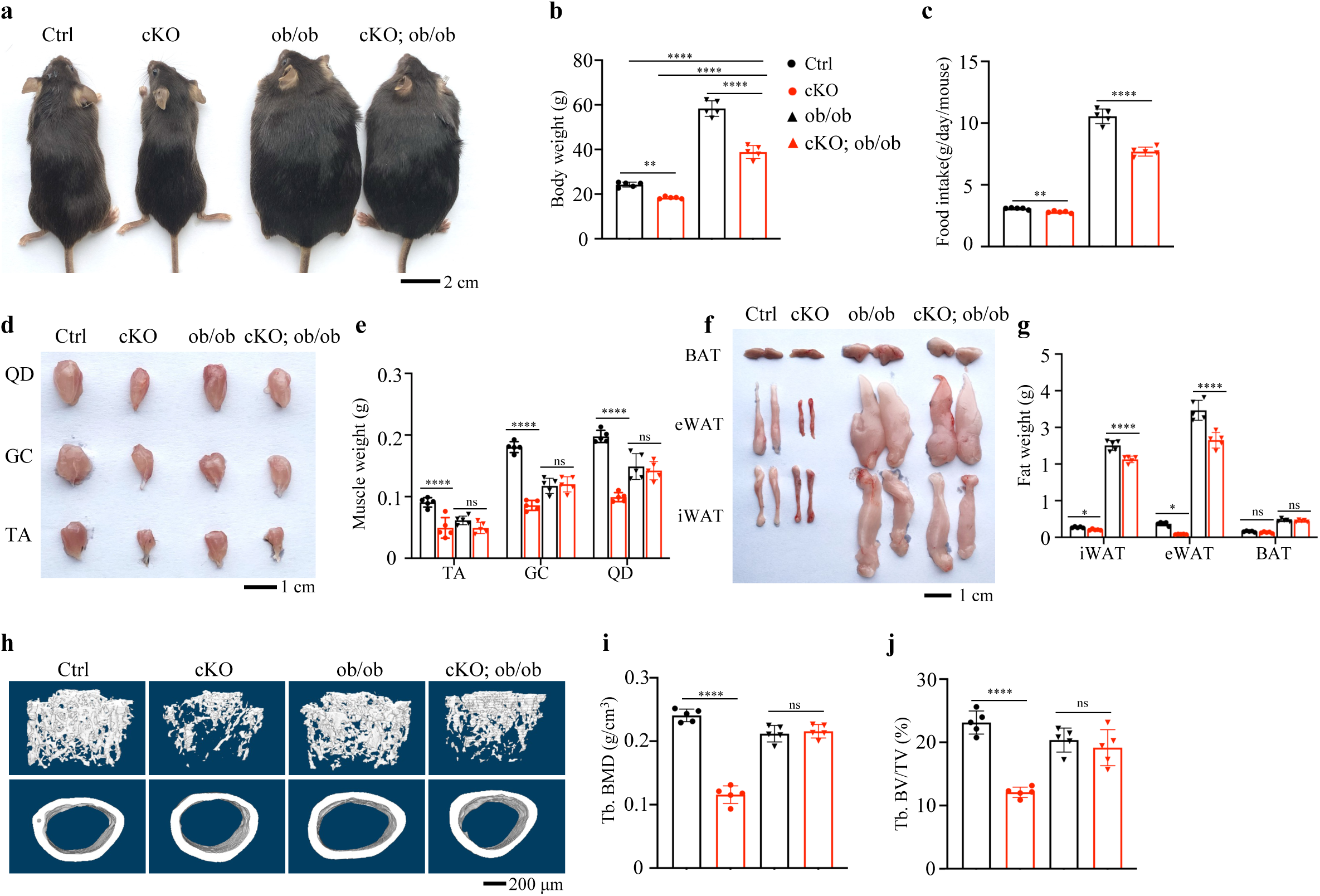
March5-dependent effects on feeding are retained in leptin-deficient mice. (**a-c**) Representative images and body weight, food intake analysis of muscle-specific March5 knockout mice on a leptin-deficient (ob/ob) background. Scale bar, 2 cm. (**d and e**) Representative images and quantification of skeletal muscle from leptin-deficient March5 cKO mice. Scale bar, 1 cm. (**f and g**) Representative images and quantification of adipose tissue from leptin-deficient March5 cKO mice. Scale bar, 1 cm. (**h-i**) Representative 3D micro-CT reconstruction and quantitative analysis of bone mass in leptin-deficient March5 cKO mice. Scale bar, 200 μm. Male mice, N = 5 per group. *P < 0.05, **P < 0.01, ***P < 0.001, ****P < 0.0001, versus controls. Student’s *t* test. Results are expressed as the mean ± standard deviation (sd).

Compared with ob/ob mice, cKO;ob/ob mice showed altered tibialis anterior, gastrocnemius and quadriceps muscle weights (Fig. 8d,e). Inguinal and epididymal white adipose tissue weights were lower in cKO;ob/ob mice than in ob/ob mice (Fig. 8f,g). Trabecular bone mineral density and bone volume fraction were also altered in cKO;ob/ob mice (Fig. 8h–j).

## Discussion

Skeletal muscle is increasingly recognized as an active regulator of organismal metabolism, particularly under conditions of aging, disuse and cellular stress. In this study, we identify March5 as a muscle proteostatic checkpoint that restrains ATF4-dependent endocrine stress signalling. March5 abundance was reduced in aged and atrophic muscle, and muscle-specific March5 deletion induced progressive post-weaning wasting characterized by reduced food intake, loss of skeletal muscle, adipose tissue and bone mass, and impaired physical performance. Conversely, muscle March5 overexpression produced reciprocal anabolic phenotypes in muscle and bone. Mechanistically, March5 associated with ATF4 and promoted ATF4 ubiquitination through a K92-dependent mechanism, thereby limiting ATF4 stability and GDF15 production. Genetic deletion of Atf4, neutralization of GDF15 and restoration of food availability each attenuated distinct components of the March5- deficient phenotype, whereas increasing March5 abundance or attenuating ATF4-associated signalling improved feeding, body composition and physical performance in aged mice. These findings define a March5–ATF4 axis that links muscle protein quality control to systemic endocrine remodelling during muscle stress and aging.

The principal mechanistic advance of this work is the identification of March5-dependent regulation of ATF4 protein stability in skeletal muscle. ATF4 is classically induced through the integrated stress response, in which phosphorylation of eIF2α promotes preferential translation of Atf4 mRNA^35^. In March5-deficient muscle and March5-depleted myogenic cells, ATF4 protein increased without a corresponding increase in Atf4 transcript abundance, indicating regulation at the post-transcriptional level. The ability of wild-type March5, but not the ligase-impaired M5^H43W^ mutant, to reduce ATF4 abundance and increase ATF4 ubiquitination places March5 E3 ligase activity upstream of ATF4 turnover. Mutational analysis further identified K92 as a critical determinant of March5-dependent ATF4 ubiquitination and degradation.

This mechanism positions March5 upstream of a known ATF4–GDF15 stress-response pathway. Prior studies have shown that mitochondrial dysfunction and integrated stress signalling can induce GDF15, including in skeletal muscle, and that muscle-derived GDF15 can remodel systemic energy balance under mitochondrial stress^21,31–33^. Therefore, the novelty of the present study is not the identification of GDF15 as an appetite-suppressive hormone. Rather, our findings define a muscle-intrinsic ubiquitin-dependent mechanism that restrains pathological activation of ATF4 and thereby limits GDF15-associated endocrine stress output. This distinction is important because it shifts the conceptual focus from GDF15 itself to the upstream proteostatic checkpoint that determines when skeletal muscle engages this endocrine programme.

The systemic phenotype of March5-deficient mice involved multiple tissues and physiological outputs. Loss of March5 in muscle reduced food intake and was accompanied by loss of lean mass, white adipose tissue, bone mass and exercise capacity. Pair-feeding experiments showed that food restriction was sufficient to reproduce a substantial portion of the body-weight and body-composition loss observed in March5-deficient mice. However, pair-fed control mice regained weight after restoration of ad libitum feeding, whereas March5-deficient mice remained lighter, indicating that reduced food intake does not fully explain the phenotype. These data support a model in which muscle-intrinsic March5 loss and secondary reduction in nutrient intake jointly contribute to systemic wasting. The incomplete rescue by Atf4 deletion and GDF15 neutralization similarly suggests that additional ATF4-dependent and ATF4-independent outputs may participate in the full phenotype.

GDF15 emerged as the most prominent endocrine factor induced by March5 deficiency. Gdf15 mRNA was strongly upregulated in March5-deficient muscle, muscle GDF15 protein was increased and circulating GDF15 concentrations were markedly elevated. Neutralization of GDF15 increased food intake and improved body weight, muscle mass, bone parameters and physical performance in March5-deficient mice. These data identify GDF15 as an important downstream effector of the March5–ATF4 axis. At the same time, GDF15 should be interpreted within its broader biological context. GDF15 can contribute to anorexia and wasting in several disease states, but it can also exert adaptive or protective effects in metabolic stress, inflammation and aging-related contexts. Thus, the March5–ATF4–GDF15 pathway may represent a context-dependent endocrine stress response rather than a uniformly detrimental signalling cascade^16,30,31,34,51,52^.

The bidirectional genetic data strengthen the functional link between March5 abundance and musculoskeletal homeostasis. March5 deletion caused loss of body mass, muscle mass and bone mass, whereas muscle March5 overexpression increased body weight, muscle mass, trabecular bone parameters and physical performance. These reciprocal phenotypes indicate that March5 abundance is functionally coupled to musculoskeletal state. Therefore, the overexpression data support bidirectional regulation of muscle and bone phenotypes, while the feeding arm of the pathway is most directly supported by the loss-of-function, GDF15 neutralization, Atf4 deletion and aged-intervention experiments.

The aging-intervention studies further indicate that the March5–ATF4 axis remains modifiable in later life. AAV-mediated March5 expression in aged mice increased food intake, body weight, skeletal muscle mass, bone mass and physical performance, and reduced circulating GDF15. Systemic ISRIB treatment produced concordant effects. These findings connect the genetic mechanism identified in March5-deficient mice with aging-associated phenotypes. Nevertheless, the intervention data should be interpreted with appropriate consideration of tissue specificity. Intravenous AAV9 delivery, even when paired with a muscle-directed promoter, may affect tissues beyond skeletal muscle, and systemic ISRIB treatment can modulate integrated stress signalling in multiple organs. Thus, these experiments establish modifiability of the March5–ATF4–GDF15 axis in aged mice.

The ob/ob cross provides an additional test of the pathway under a hyperphagic condition. Muscle March5 deletion reduced food intake and body weight in leptin-deficient mice and altered adipose, muscle and bone parameters. These results indicate that March5-dependent feeding suppression can be observed in the absence of leptin. However, because March5 deficiency causes severe wasting in non-obese mice, these data should not be interpreted as evidence that March5 inhibition is a therapeutic strategy for obesity. Instead, the ob/ob experiment supports pathway independence from leptin deficiency and extends the genetic evidence that muscle March5 status can influence feeding and body composition under distinct metabolic contexts.

In summary, this study identifies March5-dependent ATF4 ubiquitination as a skeletal muscle proteostatic mechanism that limits endocrine stress signalling. Loss of March5 stabilizes ATF4, induces a GDF15-associated endocrine programme and contributes to reduced feeding and systemic wasting. Conversely, increasing March5 abundance or attenuating ATF4-associated signalling improves aging-associated changes in feeding, body composition and physical performance in mice. These findings provide a mechanistic framework linking muscle protein quality control to organismal metabolic adaptation during muscle stress and aging.

### Limitations of the study

The human data are consistent with the mouse findings but remain supportive rather than definitive. March5 staining was reduced in skeletal muscle from older donors, whereas ATF4 and GDF15 immunoreactivity were increased. However, the human cohort was relatively small, cross-sectional and limited in available phenotypic information. In addition, circulating GDF15 can arise from multiple tissues and is influenced by renal function, inflammation, malignancy, cardiovascular disease, mitochondrial stress and other pathological conditions. Therefore, the current human data do not establish skeletal muscle as the dominant source of circulating GDF15 in older individuals. Larger cohorts with detailed assessment of physical activity, muscle mass, muscle function, nutritional state, renal function, inflammatory markers and circulating GDF15 will be required to determine the relevance of the March5–ATF4 axis to human aging and wasting disorders.

## Acknowledgment

We acknowledge the assistance of Core Research Facilities of Southern University of Science and Technology. This work was supported, in part, by the National Key R&D Program of China (2024YFA0919200), the National Natural Science Foundation of China Grants (82561160166, 82372476, 825B2074 and 824B2074), the Shenzhen Medical Research Fund (B2504003), Guangdong Provincial Science and Technology Innovation Council Grant (2017B030301018).

## Author contributions

Study design: Guixing Ma and Huiling Cao. Study conducts, data collection and analysis: Guixing Ma, Yong Chen, Siyuan Cheng, Yangshan Chen, Wei Pang, Litong Chen, and Huiling Cao. Data interpretation: Huiling Cao, Guixing Ma, Yong Chen. Drafting the manuscript: Huiling Cao and Guixing Ma. Huiling Cao takes the responsibility for the integrity of the data analysis.

## Data availability

RNA sequence and WB band raw data is stored in Mendeley Data with access number 10.17632/32jywwpk2n.1. All data generated for this study are available from the corresponding authors upon reasonable request.

## Code availability

No additional code was used in this study

## Materials and Methods

### Animal study

Genetically engineered *March5^fl/fl^*, *ATF4^fl/fl^*and *Leptin^+/-^* mice were obtained from the Shanghai Model Organisms Center, Inc. (Shanghai, China). To generate conditional knockout mice, *CKMM^Cre^* transgenic mice ^53^ (with Cre recombinase expression under the control of the CKMM promoter) were crossed with *March5^fl/fl^* mice and/or *ATF4^fl/fl^* mice. The resulting mouse models included *CKMM^Cre^; March5^fl/fl^*mice (hereafter referred to as cKO), *CKMM^Cre^; ATF4^fl/fl^*mice (aKO), *CKMM^Cre^; March5^fl/fl^; ATF4^fl/fl^*mice (dKO), *Leptin^-/-^*(ob/ob) or *CKMM^Cre^; March5^fl/fl^; Leptin^-/-^* (cKO; ob/ob). Littermates that were negative for Cre expression were used as control (Ctrl) mice. All mouse strains were maintained on a C57BL/6 background. To block the ATF4-GDF15 signaling pathway, either a GDF15-neutralizing antibody ^16,46^ (20 mg/kg) was administered via intravenous injection into 4-week-old mice once a week for 4 weeks, or ISRIB ^47^ (0.25 mg/kg) was administered daily by intraperitoneal injection to 16-month-old mice for 2 months. For overexpression of March5 specifically in skeletal muscle in mice, adeno-associated virus serotype 9 (AAV9) containing the *March5* gene driven by the ACTA1 promoter ^54,55^ was injected intravenously into 14-month-old C57BL/6 mice at a dosage of 4 × 10^12^ vector genomes (vg)/mouse. Control mice received either the solvent or empty AAV9 vectors. Mice were housed at the density of 5 per cage, with ad libitum access to food and water. The environmental conditions were controlled with a room temperature of 20-24℃. Food intake, respiratory exchange ratio (RER), and energy expenditure were measured using a comprehensive laboratory animal monitoring system (CLAMS; Columbus Instruments, USA), with each mouse housed individually in a metabolic cage. All experimental procedures involving animals were reviewed and approved by the Institutional Animal Care and Use Committee (IACUC) at the Southern University of Science and Technology, in accordance with all relevant ethical guidelines and regulations for animal research.

### Pair Feeding

Female Ctrl or cKO mice (4 weeks old) were housed in a specific pathogen-free facility at 22 ± 1 °C with a 12-hour light/dark cycle and ad libitum access to water. Mice were randomly assigned to three groups, control group (Free fed with standard chow diet), cKO group (cKO mice free fed with standard chow diet), Pair-fed group (cKO mice Fed isocaloric standard chow diet matched daily to the mean intake of the cKO group).

### Pulse oximetry

Peripheral blood oxygen saturation (SpO₂) was repeatedly measured using a mouse pulse oximeter equipped with a MouseOx collar sensor (Starr Life Sciences Corp., USA). Hair around the neck and shoulder area, where the collar sensor was placed, was carefully removed before measurement. SpO₂ was continuously recorded for more than 3 minutes. The mean value of stable, artifact-free readings was used to determine the SpO₂ for each mouse^56^.

### Human samples

Human skeletal muscle samples were obtained from participants at Nanjing Drum Tower Hospital (Nanjing, China) following procedures approved by the hospital’s ethics committee (IRB No: 2021-192). Written informed consent was obtained from all participants prior to sample collection. Detailed information for both young and aged individuals is provided in Supplementary Table 2. Participates with other metabolic disorders, such as diabetes, arthritis, etc., were excluded from this study to minimize confounding factors.

### Cell culture

C2C12 myoblasts were cultured in Dulbecco’s Modified Eagle Medium (DMEM) supplemented with 10% fetal bovine serum and 1% penicillin-streptomycin at 37 °C in a humidified incubator with 5% CO_2_. To induce myogenic differentiation, upon reaching 80–90% confluence, the cells were switched to differentiation medium containing 2% horse serum and 1% penicillin-streptomycin for 5 days to promote myotube formation. For gene silencing, C2C12 cells were transfected with small interfering RNA targeting March5 (siMarch5, referred to as siM5) following the manufacturer’s protocol using Lipofectamine RNAiMAX (Invitrogen, #13778100). Details of siM5 and the negative control siRNA (siNC) are provided in Supplementary Table 3. Transfection of a March5 overexpression plasmid (M5) was performed according to the manufacturer’s instructions using Lipofectamine 3000 (Invitrogen, # L3000015), as previously described ^57^.

### Micro-computerized tomography (μCT) analysis

Fixed femurs were used for μCT analysis using a Bruker μCT (SkyScan 1172 Micro-CT, Bruker MicroCT), as previously described ^58^. The scanning procedure adhered to the standards and terminology recommended by the American Society for Bone and Mineral Research (ASBMR). Briefly, non-demineralized femurs were scanned at 60 KV, 100 μA, and an exposure time of 926 ms, with a voxel size of 10 μm. For trabecular bone analysis, a region of interest (ROI) with a length of 1.5 mm, starting from 0.5 mm proximal to the distal growth plate, was selected. For cortical bone analysis, a 1.0 mm ROI at the mid-diaphysis of the femur was analyzed. The following bone parameters were measured: bone mineral density (BMD), bone volume fraction (BV/TV), and cortical thickness (Cort.Th.).

### Histological evaluation

Tissue samples were fixed in 4% paraformaldehyde (PFA) overnight at 4 ℃ ^59^, followed by dehydration and paraffin embedding. Paraffin-embedded sections, 5-μm in thickness, were subsequently deparaffinized, rehydrated, and used for hematoxylin and eosin (HE), immunohistochemistry (IHC), and immunofluorescence (IF) staining.

### Real-time quantitative reverse transcription PCR (qRT-PCR) analyses

RNA isolation, reverse transcription, and qRT-PCR analyses were performed as previously described ^60^. Total RNA was extracted using Trizol reagent and further purified with a total RNA isolation kit (Transgen, ER601-01). The RNA quality and concentration were assessed by measuring the 260/280 ratio using a NanoDrop 3000 spectrophotometer. Subsequently, 1 μg of mRNA was reverse transcribed into complementary DNA (cDNA) using a reverse transcription reagent kit (Beyotime, D7190). Quantitative PCR amplification was performed using SYBR Green Master Mix (Beyotime, D7260) ^61^. Gene expression levels were calculated using the ΔCt-method. The mRNA primers were synthesized by Sangon Biotech, and the sequences of the primer pairs used in this study are provided in Supplementary Table 4.

### Western blot (WB) analysis

Protein isolation and Western blot analyses were conducted as previously described ^60^. Cells were lysed in RIPA buffer (Sigma, R0278), and the protein concentration was determined using a BCA protein assay kit (Cwbio, CW0014). Equal amounts of protein lysates were separated by SDS-PAGE and subsequently transferred onto polyvinylidene difluoride (PVDF) membranes. Membranes were incubated with primary antibodies at 4 ℃ for 12 hours, followed by incubation with horseradish peroxidase (HRP)-conjugated secondary antibodies. Detailed information on the antibodies used is provided in Supplementary Table 5 Protein bands were visualized using a chemiluminescent HRP substrate. Quantification of Western blot results was performed using ImageJ software ^62^. In brief, the grayscale values of the protein bands were analyzed, and the expression of target proteins was normalized to internal control proteins for quantification.

### Immunohistochemistry (IHC) and immunofluorescence (IF) staining

Tissues were dehydrated and embedded in paraffin. Paraffin-embedded sections, 5-μm in thickness, were deparaffinized and subjected to antigen retrieval in citric acid buffer at 58 ℃ for 16 hours. For IHC staining, 5 μm sections were incubated with the specified primary antibodies or control IgG using the EnVision+System-HRP (DAB) kit (Dako North America Inc, K401111-2), according to the manufacturer’s instruction ^63^. Images were acquired using a Lecia Aperio VERSA scanner. For IF staining, 5-μm sections were incubated with primary antibodies at 4 ℃ for 8 hours, followed by incubation with fluorescent-labeled secondary antibodies for 1 hour at room temperature. Confocal images were captured using Zeiss LSM980 confocal microscope. Quantification of staining was performed using ImageJ software, with all data normalized to the control group.

### Co-immunoprecipitation (Co-IP) assay

The co-immunoprecipitation (Co-IP) assay was performed as previously described ^63^. Briefly, cells were lysed in immunoprecipitation (IP) lysis buffer (pH 7.4, 0.025 M Tris, 0.15 M NaCl, 0.001 M EDTA, 1% NP-40, and 5% Glycerol) (Thermo Fisher), supplemented with a protease inhibitor cocktail. The lysates were incubated on ice for 10 minutes, followed by centrifugation at 13,000 × g for 10 minutes at 4 °C. The resulting supernatant was incubated overnight at 4°C with the appropriate primary antibody, followed by 1 hour incubation at room temperature with Protein A/G magnetic beads. The antigen-antibody-bead complex was isolated using a DynaMag™-2 Magnet (Thermo Fisher) and washed three times with IP lysis buffer. The bound protein complexes were eluted by resuspending the beads in 60 μl of 1× loading buffer and heating at 95 °C for 5 minutes. The samples were then resolved by SDS-PAGE and analyzed by Western blotting.

### Structure prediction

The structural prediction of the March5-ATF4 complex was performed using AlphaFold-Multimer ^49^, following previously established protocols. The resulting structural models were visualized and analyzed using Pymol and Chimera software ^64^.

### Quantification and statistical analysis

Mice were randomly assigned to experimental groups. IF, IHC, and histological analyses were performed and evaluated in a double-blinded manner. Statistical analyses were conducted using GraphPad Prism 9 (Version 9.5.0). Data are presented as the mean ± standard deviations (sd). Statistical significance was assessed using either a two-tailed Student’s t-test or one-way ANOVA, depending on the comparison being made. Survival analysis was performed using the Log-rank test. p < 0.05 was considered statistically significant.

## Supplementary Tables

**Table S1.** March5-related diseases showed in Genebass Database.

| Description | N cases | N controls | P-Value | Beta |
| --- | --- | --- | --- | --- |
| W45.1 Primary total prosthetic replacement of joint NEC | 264 | 394577 | 0.00043423 | 0.24319665 |
| T72.3 Release of constriction of sheath of tendon | 397 | 394444 | 0.00099957 | 0.21575158 |
| Lansoprazole | 14214 | 380569 | 0.00126049 | 0.12093355 |
| Vitamin b6 preparation | 399 | 213189 | 0.00154785 | 0.2009991 |
| M66 Spontaneous rupture of synovium and tendon | 1014 | 393827 | 0.0016364 | 0.19356282 |
| **Z56.9 Muscle of hand NEC** | **726** | **394115** | **0.00177255** | **0.19171115** |
| K51.2 Intravascular ultrasound of coronary artery | 152 | 394689 | 0.00201332 | 0.37614582 |
| total fatty acids | 369 | 394414 | 0.003666355 | 0.318205729 |
| Hodgkins lymphoma / hodgkins disease | 1208 | 393575 | 0.004064918 | 0.161232937 |
| Selenium product | 362 | 213251 | 0.004408653 | 0.31511818 |

**Table S2.**
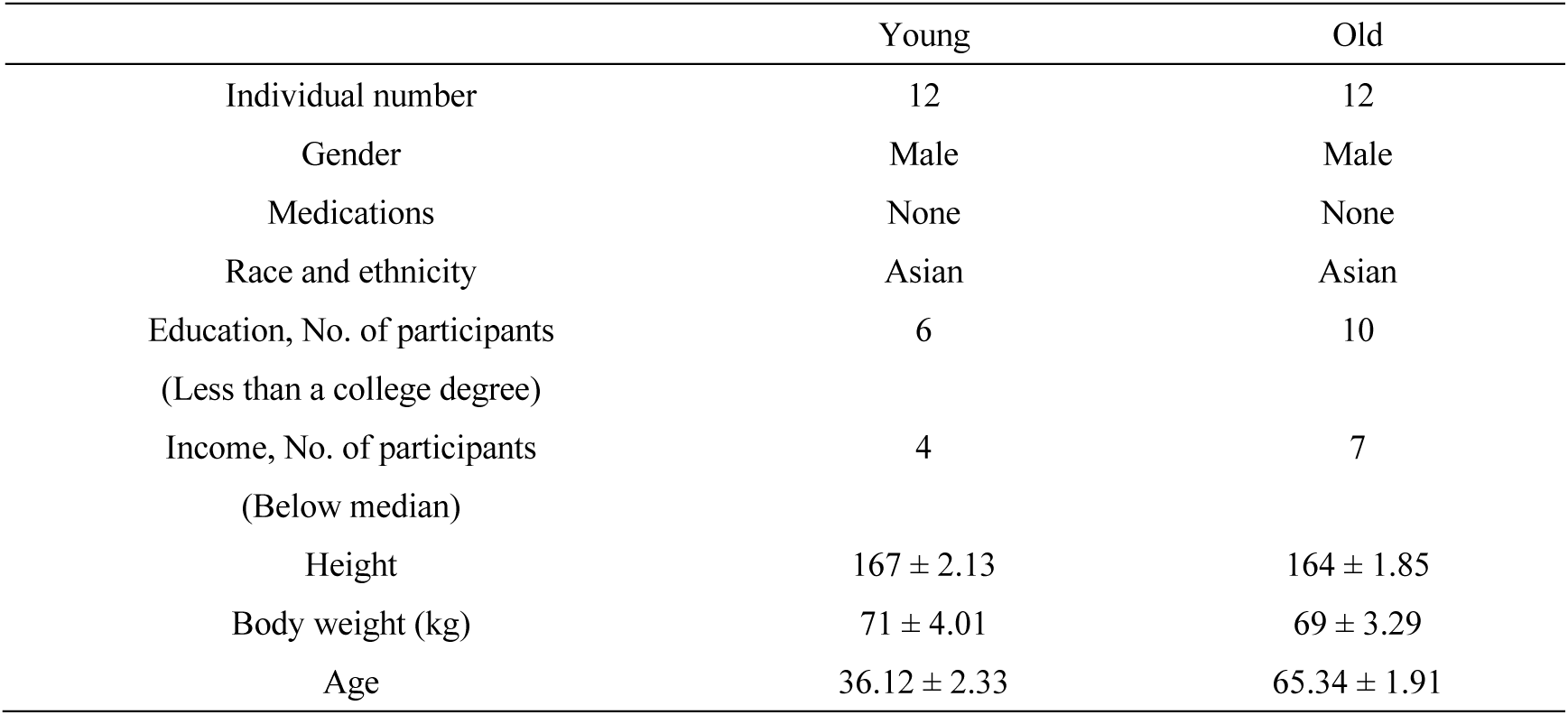
Demographics of the human individuals involved in this study.

|  | Young | Old |
| --- | --- | --- |
| Individual number | 12 | 12 |
| Gender | Male | Male |
| Medications | None | None |
| Race and ethnicity | Asian | Asian |
| Education, No. of participants<br>(Less than a college degree) | 6 | 10 |
| Income, No. of participants<br>(Below median) | 4 | 7 |
| Height | 167 ± 2.13 | 164 ± 1.85 |
| Body weight (kg) | 71 ± 4.01 | 69 ± 3.29 |
| Age | 36.12 ± 2.33 | 65.34 ± 1.91 |

**Table S3.**
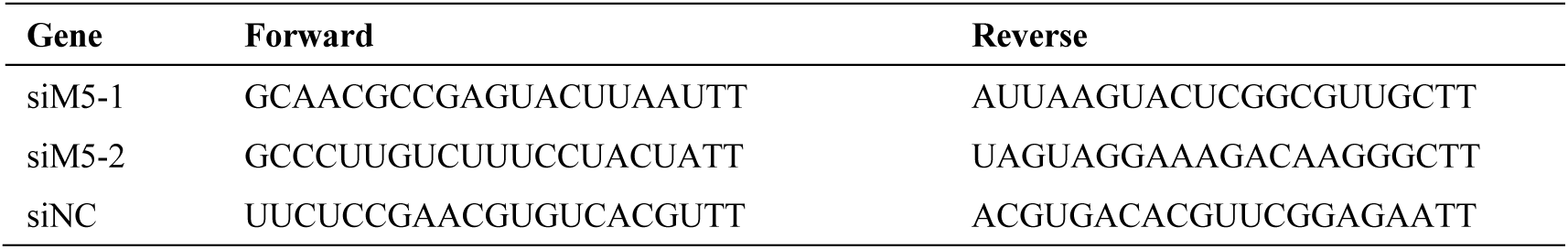
Sequence of DNA or RNA used in this study.

**Table S4.**
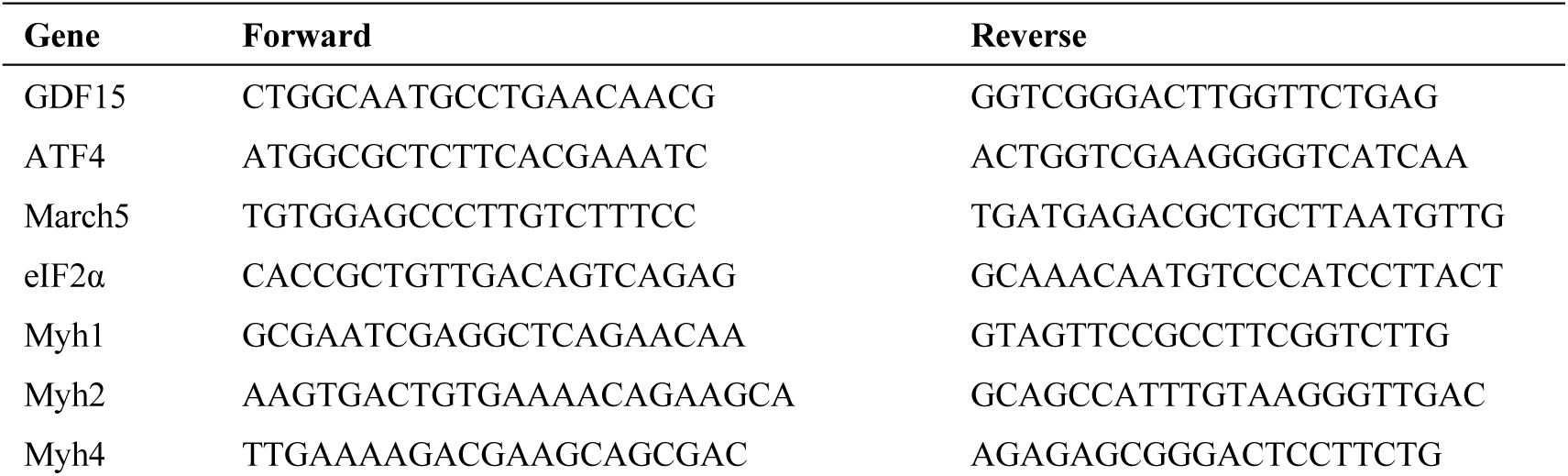
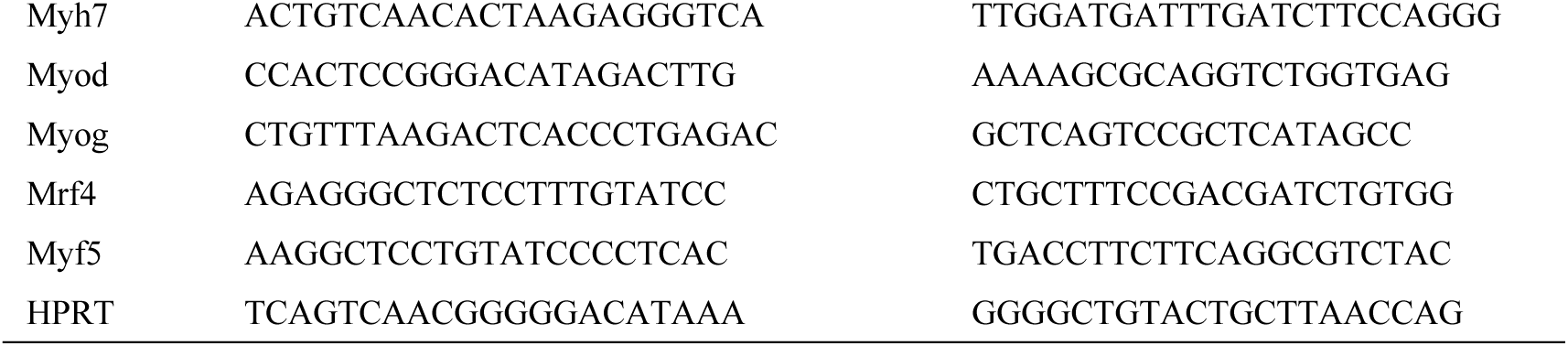
Primer information used in this study.

**Table S5.** Antibodies, chemicals, peptides, recombinant proteins and kits used in this study.

| **Antibodies** |  |  |
| --- | --- | --- |
| MYH1 | Servicebio | Cat#GB112130-100 |
| MYH7 | Servicebio | Cat#GB111857-100 |
| March5 | Absin | Cat#abs113478 |
| ATF4 | Selleck | Cat# A5514 |
| eIF2α | Abclonal | Cat#A21221 |
| p-eIF2α | Abclonal | Cat#AP0745 |
| GDF15 | Beyotime | Cat# AF6975 |
| Anti-GDF15 neutralizing antibody | DIMABio | Cat# DMC100474 |
| Actin | ZSGB-BIO | Cat# TA-09 |
| GFP | TechiSun | Cat# TAG10044 |
| UB | ABClonal | Cat# A19686 |
| **Chemicals, peptides, and recombinant proteins** |  |  |
| H<sub>2</sub>O<sub>2</sub> | Shanghai ZEYE bio | Cat#ZY65553PD |
| IL1-β | ABClonal | Cat# RP00002 |
| Lipofectamine RNAiMAX | Invitrogen | Cat#13778150 |
| PEI | Servicebio | Cat# G1805 |
| Giemsa | Sigma | Cat#GS500 |
| SYBR | Beyotime | Cat#D7260 |
| **Critical commercial assays** |  |  |
| Osteo-Bed Bone Embedding kit | Sigma | Cat#EM0T48598200 |
| GDF15 ELISA kit | TechiSun | Cat#TQ48598 |
| Total RNA isolating kit | Transgen | Cat#ER601-01 |
| First-standard cDNA synthesis kit | GeneCopeia | Cat#QP114 |
| miRNA RT-qPCR detection kit | GeneCopeia | Cat#QP011 |
| EnVision+System-HRP (DAB) kit | Dako | Cat#K401111-2 |

## Extended Data Figures

**Extended Data Figures 1.**
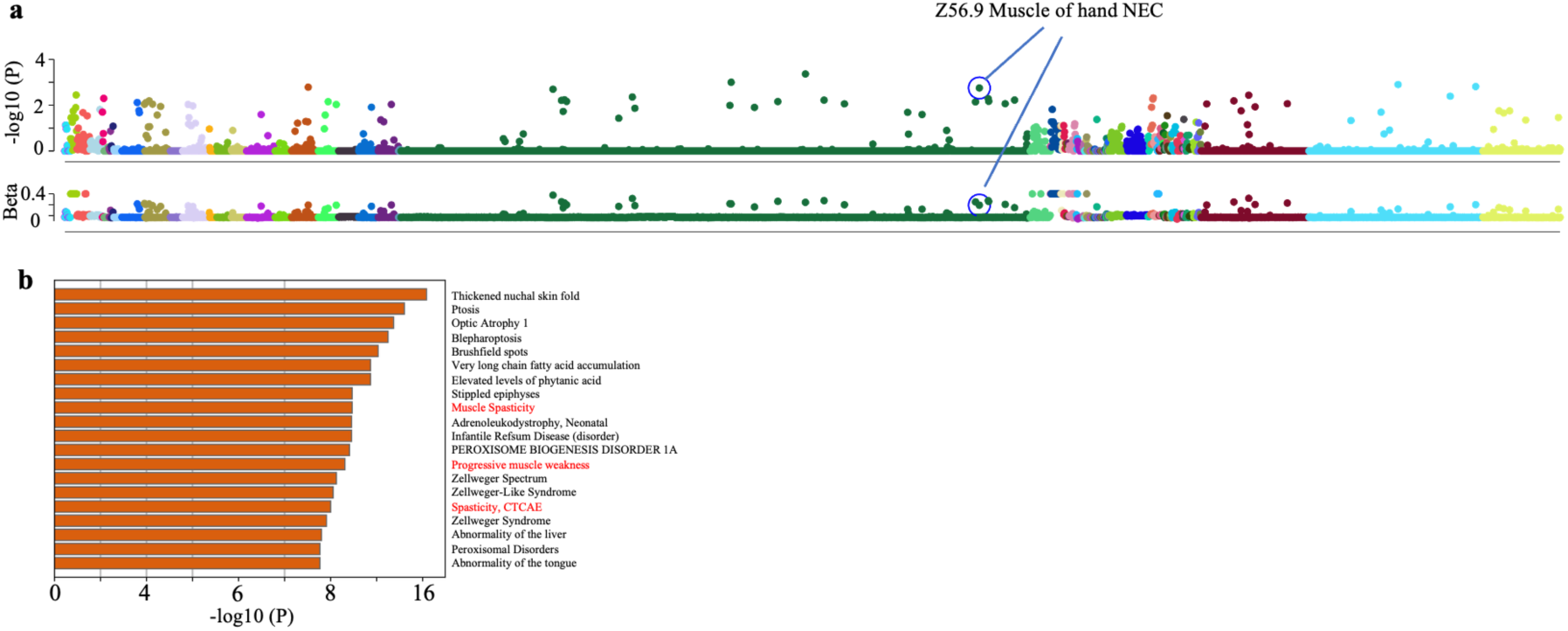
March5 is associated with muscle function. (**a**) Heatmap displaying associated diseases with the March5 interactome. (**b**) Manhattan plot illustrating March5-related diseases based on data from Genebass Database.

**Extended Data Figures 2.**
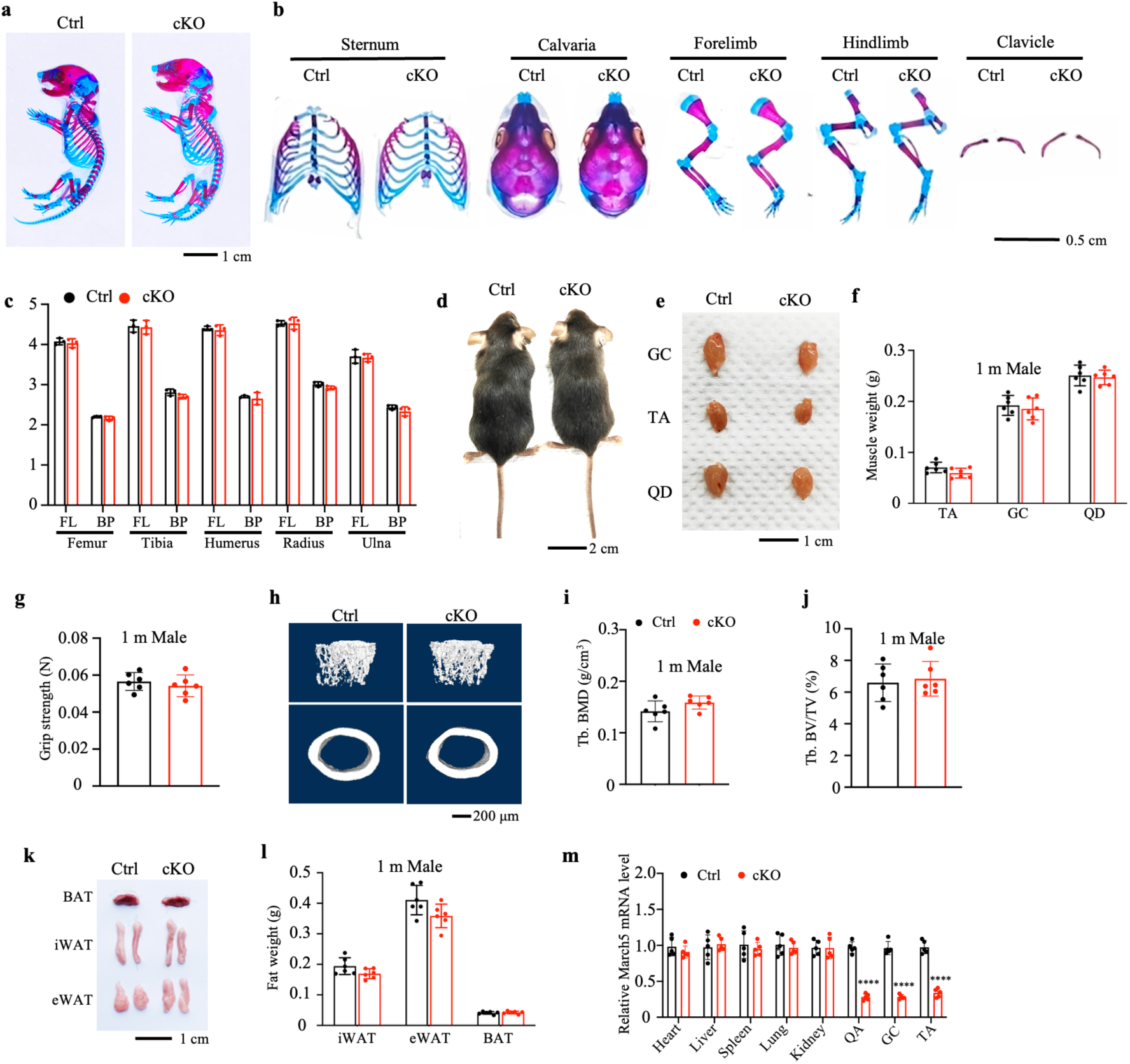
Skeletal muscle-specific March5 knockout does not affect musculoskeletal development. (**a**) Alizarin red and Alcian blue double staining of skeletons of postnatal day 0 (P0) cKO and littermate control mice. (**b and c**) Representative images and quantitative analysis of the full-length (FL) and bony portion (BP) length of the sternum, calvaria, forelimb, hindlimb, and clavicle. Male mice, N = 3 per group. (**d and e**) Representative images of gastrocnemius (GC), tibialis anterior (TA), and quadriceps (QD) muscles in cKO and control mice. Scale bar, 2 cm (**d)**, 1 cm (**e)**. (**f**) Quantification of muscle size in (**e).** (**g**) Quantitative analyses of forelimb grip strength. (**h-j**) 3D reconstruction (**h**) and quantitative analysis of BMD (**i**) and BV/TV (**j**) from μCT scans of the distal femurs of cKO and control mice. Scale bar, 200μm. Male mice, N = 6 per group. (**k**) Representative images of BAT, iWAT, eWAT in cKO and littermate control mice. (**l**) Quantification of adipose tissue weight from cKO and littermate control mice (**k**). (**m**) qRT-PCR analyses of RNA samples isolated from organs of cKO and littermate control mice. Scale bar, 1 cm. Male mice, N = 6 per group. *P < 0.05, **P < 0.01, ***P < 0.001, ****P < 0.0001, versus controls. Student’s *t* test. Results are expressed as the mean ± standard deviation (sd).

**Extended Data Figures 3.**
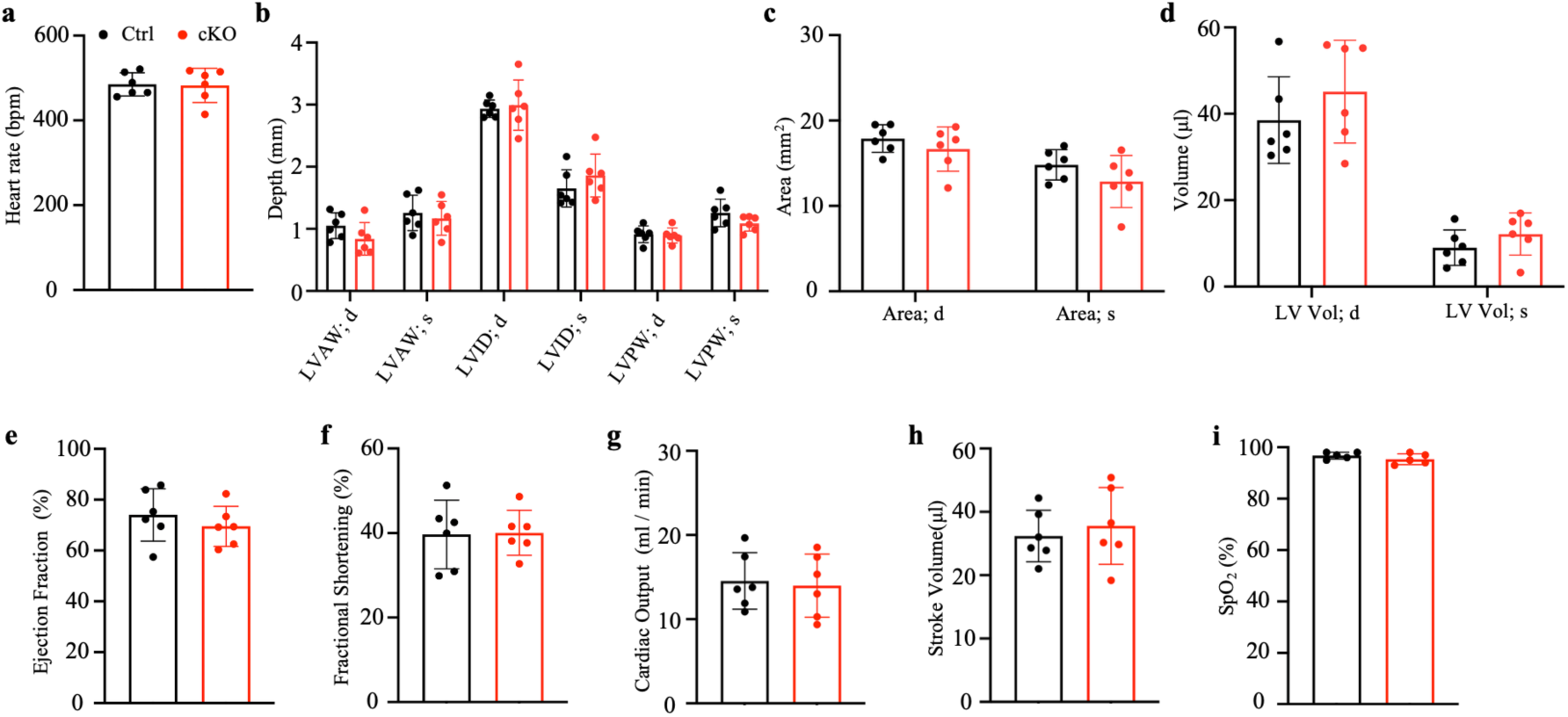
Skeletal muscle-specific March5 knockout does not affect heart function. (**a-d**) Statistics of heart rate (beats per minute, bpm) (**a**); left ventricle anterior wall thickness during diastole (LVAW; d) and systole (LVAW; s), left ventricular internal dimension during diastole (LVID; d) and systole (LVID; s), left ventricle posterior wall thickness during diastole (LVPW; d) and systole (LVPW; s) (**b**); end-diastolic (Area; d) and end-systolic (Area; s) ventricular area (**c**); and left ventricular end-diastolic volume (LV Vol; d) and end-systolic volume (LV Vol; s) (**d**) in cKO and littermate control mice at 2 months of age. Male mice, N = 6 per group. (**e-h**) Statistical analysis of ejection fraction (**e**), fractional shortening (**f**), cardiac output (**g**), and stroke volume (**h**) in cKO and littermate control mice at 2 months of age. (**i**) Statistical analysis of Peripheral blood oxygen content (SpO_2_) in cKO and littermate control mice at 2 months of age. Male mice, N = 6 per group. *P < 0.05, **P < 0.01, ***P < 0.001, ****P < 0.0001, versus controls. Student’s *t* test. Results are expressed as the mean ± standard deviation (sd).

**Extended Data Figures 4.**
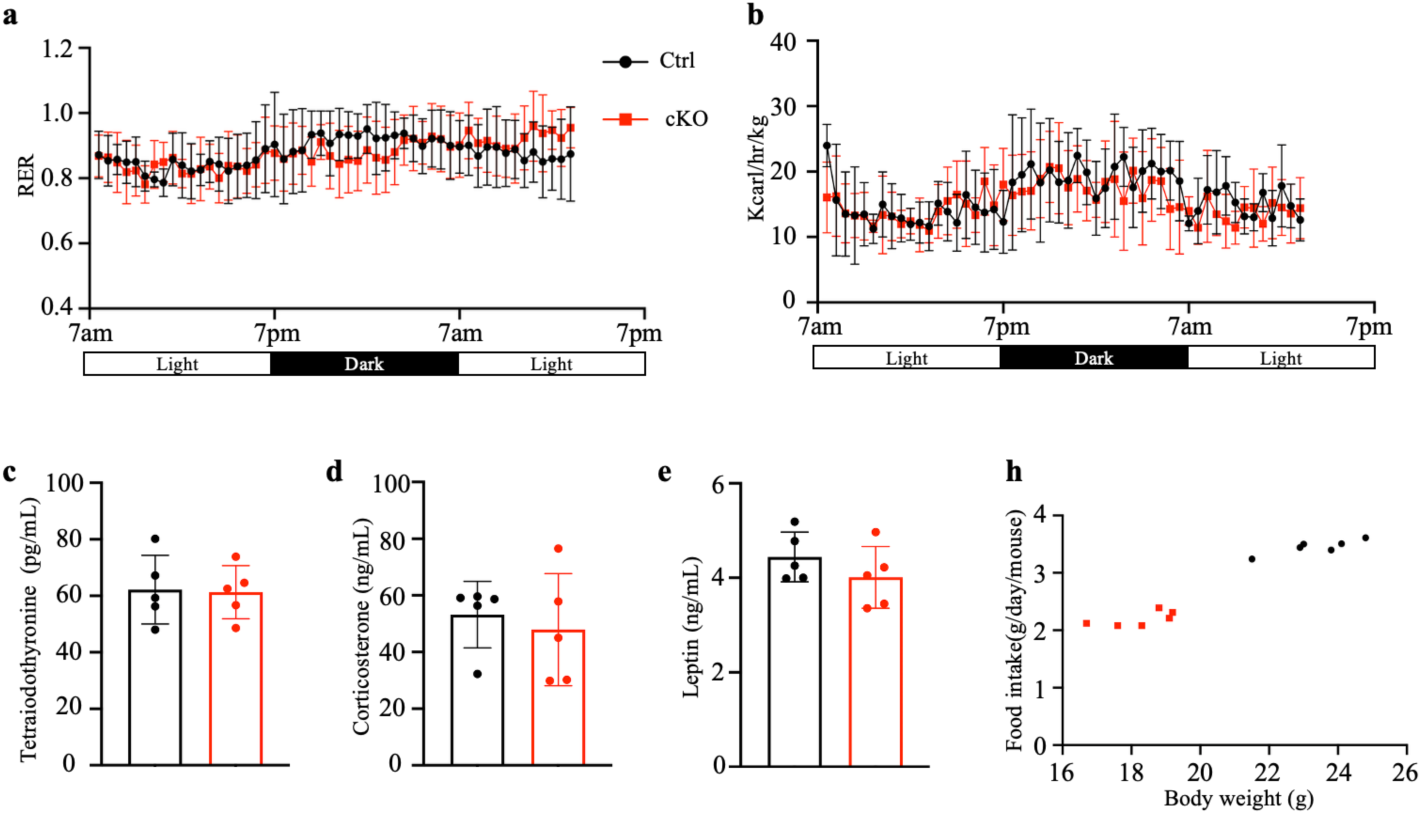
Skeletal muscle–specific deletion of March5 does not affect systemic metabolism in mice. (**a-b**) Metabolic analysis of respiratory exchange ratio (RER) and energy expenditure in control and muscle-specific March5 knockout (cKO) mice. (**c-e**) ELISA measurement of serum thyroxine (T4), corticosterone, and leptin levels in control and cKO mice. (**h**) Scatter plot showing the correlation between body weight and food intake in individual mice. Female mice, N = 5 per group. *P < 0.05, **P < 0.01, ***P < 0.001, ****P < 0.0001, versus controls. Student’s *t* test. Results are expressed as the mean ± standard deviation (sd).

**Extended Data Figures 5.**
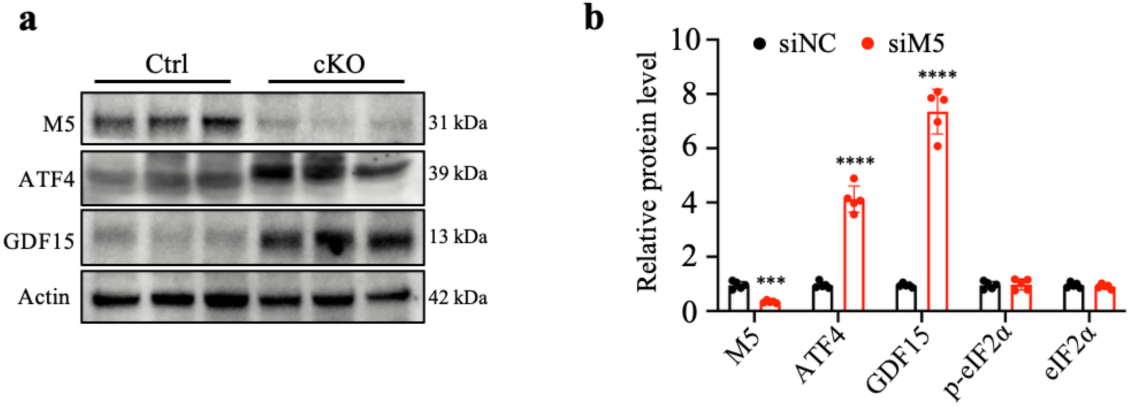
Muscle-specific deletion of March5 upregulates ATF4 and its downstream target GDF15. (**a-b**) Western blot analysis and quantification of ATF4 and GDF15 protein levels in QC lysates from Ctrl and cKO mice. Male mice, N = 5 per group. *P < 0.05, **P < 0.01, ***P < 0.001, ****P < 0.0001, versus controls. Student’s *t* test. Results are expressed as the mean ± standard deviation (sd).

**Extended Data Figures 6.**
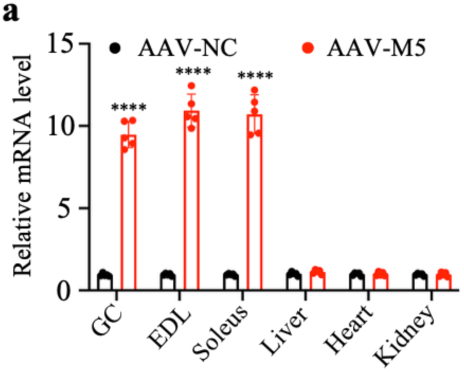
AAV-M5 efficiently targets skeletal muscle to overexpress March5. (**a**) qRT-PCR analysis of March5 mRNA levels in various tissues following AAV-M5 administration. Male mice, N = 5 per group. *P < 0.05, **P < 0.01, ***P < 0.001, ****P < 0.0001, versus controls. Student’s *t* test. Results are expressed as the mean ± standard deviation (sd).

**Extended Data Figures 7.**
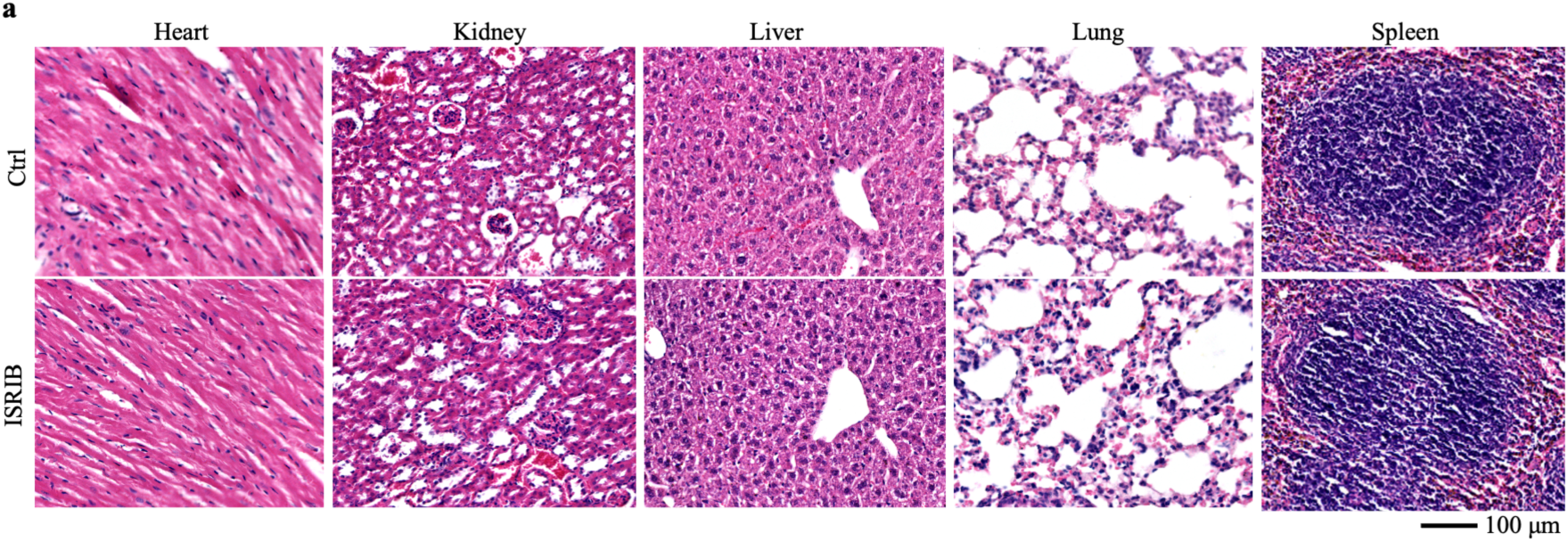
ISRIB has no toxic effect on tissues and organs *in vivo*. (**a**) Representative HE staining photographs of heart, liver, spleen, lung and kidney of mice after treatment with ISRIB or Control group. Scale bar, 100 μm.

